# Unified neural pathways that gate affective pain and multisensory innate threat signals to the amygdala

**DOI:** 10.1101/2020.11.17.385104

**Authors:** Sukjae Joshua Kang, Shijia Liu, Mao Ye, Dong-Il Kim, Jong-Hyun Kim, Tae Gyu Oh, Jiahang Peng, Ronald M. Evans, Kuo-Fen Lee, Martyn Goulding, Sung Han

## Abstract

Perception of aversive sensory stimuli such as pain and innate threat cues is essential for animal survival. The amygdala is critical for aversive sensory perception, and it has been suggested that multiple parallel pathways independently relay aversive cues from each sensory modality to the amygdala. However, a convergent pathway that relays multisensory aversive cues to the amygdala has not been identified. Here, we report that neurons expressing calcitonin gene-related peptide (CGRP) in the parvocellular subparafasicular thalamic nucleus (SPFp) are necessary and sufficient for affective-motivational pain perception by forming a spino-thalamo-amygdaloid pain pathway. In addition, we find that this thalamic CGRP pain pathway, together with well-known parabrachio-amygdaloid CGRP pain pathway, is critical for the perception of multisensory innate threat cues. The discovery of unified pathways that collectively gate aversive sensory stimuli from all sensory modalities may provide critical circuit-based insights for developing therapeutic interventions for affective pain- and innate fear-related disorders.

## Introduction

Pain is a complex sensory and emotional experience caused by tissue-damaging noxious stimuli that produce immediate avoidance behavior, as well as long-lasting aversive memories so that future damage can be avoided (Julius and Basbaum, 2001; Melzack and Casey, 1968). Therefore, the perception of pain results in behavioral outcomes similar to those associated with the perception of threats. Indeed, most research on Pavlovian threat learning has used electric foot shock, a painful noxious stimulus, as a threat cue. It has also been suggested that pain and threat perceptions interact with each other (Elman and Borsook, 2018). Individuals with pain asymbolia, who have deficits in perceiving affective-motivational aspects of pain due to damage to limbic structures, show compromised perception of threats (Berthier et al., 1988; Rubins and Friedman, 1948). Alternatively, people with affective pain disorders, such as migraine and fibromyalgia, are often hypersensitive to sensory inputs and perceive normal sensory signals as aversive (Bar-Shalita et al., 2019; López-Solà et al., 2017). Therefore, it is likely that there are unified neural circuits and brain areas that process both pain-causing noxious stimuli and threat-producing aversive sensory cues (Price, 1999).

The amygdala is a key limbic structure that integrates sensory stimuli with an internal state to generate appropriate emotional responses (Janak and Tye, 2015; LeDoux, 2012, 2000). It is activated by aversive sensory stimuli, including noxious stimuli (Ren and Neugebauer, 2010; Simons et al., 2014; Veinante et al., 2013), and lesioning the amygdala greatly attenuates the perception of multimodal sensory threats (Bach et al., 2015; Blanchard and Blanchard, 1972; Dal Monte et al., 2015) and pain (Gao et al., 2004; Helmstetter, 1992; Manning and Mayer, 1995; Tanimoto et al., 2003). Therefore, the amygdala may serve as a pivotal node in integrating and unifying all threat cues from different sensory modalities, including pain-causing noxious stimuli. Recent studies suggest that aversive sensory stimuli from each sensory modality relay threat cues to the distinct brain areas including the hypothalamus and amygdala through parallel non-overlapping pathways. These include somatosensory (Barsy et al., 2020; Choi et al., 2020; Han et al., 2015; Sato et al., 2015), visual (Salay et al., 2018; Wei et al., 2015; Zhou et al., 2019), auditory (Barsy et al., 2020), gustatory (Carter et al., 2013; Kim et al., 2017; Wang et al., 2018) and olfactory (Rosen et al., 2015; Tong et al., 2020). However, little is known about the convergent neural circuits that relay and integrate multimodal aversive sensory signals, including nociceptive stimuli, to the amygdala.

Noxious stimuli from the periphery are relayed to the brain through two ascending pain pathways, the spino-parabrachial pathway and the spino-thalamic pathway (Bushnell et al., 2013). It is a well-established idea that the spino-thalamic pathway is involved in sensory and discriminative pain perception and that the spino-parabrachial pathway is involved in the perception of affective and motivational pain. This is because the former projects to the somatosensory cortex and the latter projects to the amygdala (Basbaum et al., 2009). Among multiple areas of the amygdala, the capsular subdivision of the central nucleus of the amygdala (CeC) is known as the nociceptive amygdala since it is activated by nociceptive stimuli and receives direct input from the parabrachial nucleus (PBN) through the spino-parabrachial pathway (Gauriau and Bernard, 2002; Neugebauer, 2015). Nevertheless, it remains unclear how the nociceptive information is relayed to other areas of the amygdala involved in pain processing, such as the amygdala-striatum transition area (AStr) (Xiu et al., 2014) and the lateral nucleus of the amygdala (LA) (Bernard et al., 1992; Thompson and Neugebauer, 2017). In particular, the LA is critically involved in pain-induced aversive learning (LeDoux, 2007; Ressler and Maren, 2019), but the detailed pain pathway that relays nociceptive information to the LA has not been fully understood.

Although the thalamus has been implicated in sensory and discriminative pain (Dado et al., 1994; Zhang and Giesler, 2005), some thalamic nuclei, such as the ventromedial posterior thalamus (VMpo) in primates, or the triangular subdivision of the posterior thalamus (PoT) in rodents, are thought to be involved in affective and motivational pain perception by relaying nociceptive signals to the limbic areas (Craig et al., 1994, 2000; Gauriau and Bernard, 2004a; Price, 2002). However, the involvement of these nuclei in affective and motivational pain perception is still inconclusive (Willis et al., 2002), mainly because the thalamus has many small, functionally distinct nuclei without clear anatomical boundaries. Therefore, to understand the roles of the thalamus in affective pain processing, it is critical to identify genetically defined populations of thalamic neurons that directly convey spinal nociceptive input to the limbic areas, such as the amygdala.

Calcitonin gene-related peptide (CGRP) is a 37-amino acid neuropeptide produced by peripheral neurons and mediates vasodilation and nociception (Russell et al., 2014; Russo, 2015). It is also produced in the brain and plays an essential role in aversive learning and pain perception (Palmiter, 2018; Shinohara et al., 2017; Yu et al., 2009). CGRP-expressing neurons are highly clustered in two brain areas: the external lateral subdivision of the PBN (PBel) and the parvocellular subparafascicular nucleus (SPFp) (D’Hanis et al., 2007; Dobolyi et al., 2005); (Experiment 79587715, Allen Brain Atlas). Previous studies have shown that CGRP neurons in the PBel (CGRP^PBel^) are critically involved in transmitting affective pain signals during aversive learning (Han et al., 2015) and in transmitting visceral aversive signals to the CeA (Chen et al., 2018). On the other hand, the latter is a relatively unexplored area. The SPFp is an elongated structure that extends from the anteromedial to posterolateral thalamus (D’Hanis et al., 2007). It has been speculated that CGRP neurons in the medial part of the SPFp (CGRP^SPFp^) may play a role in sexual behaviors (Coolen et al., 2003a, 2003b), whereas those in the posterolateral part of the CGRP^SPFp^ may be involved in emotional behaviors, based on anatomical projections to the amygdala (D’Hanis et al., 2007; LeDoux et al., 1985; Yasui et al., 1991). The neighboring thalamic nucleus, such as the posterior intralaminar nucleus (PIL), or medial subdivision of medial geniculate nucleus (MGm) have been implicated to relay noxious stimuli to the amygdala during aversive learning (Barsy et al., 2020; Shi and Davis, 1999). However, it has not been tested whether the CGRP^SPFp^ neurons are involved in conveying pain signals to the amygdala during affective-motivational pain perception. Furthermore, it is unknown to what extent CGRP^PBel^ and CGRP^SPFp^ neurons play similar roles in conveying aversive sensory information to the amygdala.

Here, we report that CGRP^SPFp^ neurons receive direct monosynaptic inputs from projection neurons within the dorsal horn of the spinal cord and project their axons to multiple regions within the amygdala (namely (the AStr and the LA) and to the posterior insular cortex, but not to the somatosensory cortex. These neurons are activated by multimodal nociceptive stimuli. Silencing these neurons substantially attenuates affective and motivational pain perception, and activating these neurons induces aversion and aversive memory. Furthermore, CGRP^SPFp^ neurons, together with CGRP^PBel^ neurons, are collectively activated by aversive sensory stimuli from all sensory modalities (visual, auditory, somatosensory, gustatory, and olfactory), and silencing these neurons attenuates the perception of all aversive sensory stimuli. Taken together, CGRP neurons within the SPFp and PBel not only form two affective pain pathways, namely the spino-thalamo-amygdaloid and the spino-parabracho-amygdaloid pathways, but they also convey aversive sensory signals from all sensory modalities to the amygdala during threat perception.

## Results

### CGRP^SPF^p and CGRP^PBel^ neurons connect multiple sensory relay areas to the amygdala

The amygdala is critically involved in the affective-motivational pain perception. However, it is not fully understood by which the nociceptive information is conveyed to the amygdala. The spinal dorsal horn projection neurons relay nociceptive signals to the brain (Basbaum et al., 2009; Todd, 2010). One example is *Tacr1*-expressing neurons in the spinal dorsal horn (Barik et al., 2020; Chiang et al., 2020; Choi et al., 2020; Deng et al., 2020). To identify direct spino-recipient areas in the brain, we genetically labeled only the spinal *Tacr1* neurons with tdTomato fluorescent protein by the triple crossing of *Tacr1^Cre^*, *Cdx2^FlpO^*, and *Ai65* (Rosa-CAG-FrtSTOPFrt-LoxSTOPLox-tdTomato; *dsTomato*) mice as described previously (Bourane et al., 2015) (Figure S1A). The tdTomato-expressing cell bodies were only observed in the spinal dorsal horn (Figure S1A). Fluorescently labeled axonal terminals were observed in multiple brain areas, including the PBN, SPFp, the posterior complex of the thalamus (Po), the ventral posterolateral nucleus of the thalamus (VPL), superior colliculus (SC), periaqueductal gray (PAG), dorsal column nuclei (DCN), and ventrolateral medulla (VLM) (Figure S1B). We then asked which of these areas project to the amygdala. By searching through the Allen Mouse Connectivity Atlas (http://connectivity.brain-map.org/), we found that the SPFp and PBel project to the LA and CeA, respectively. Interestingly, CGRP neurons are found in both the SPFp and PBel, and CGRP^PBel^ neurons are known to play a role in affective pain perception (Han et al., 2015). Therefore, we sought to dissect and compare the roles of these CGRP circuits in processing nociceptive sensory information and relaying this information to the amygdala.

To identify regions that lie downstream of CGRP neurons in the SPFp and PBel, we injected Cre-dependent AAVs encoding EYFP or mCherry into the SPFp and PBel of the same *Calca^Cre^* mouse that expresses Cre-recombinase in the CGRP-expressing neurons (*Calca* gene encodes CGRP), respectively (Figure 1A). Coronal slices around AP −1.1 showed an intermingled green and red expression pattern in the CeA and LA (Figure S2A). However, posterior slices (AP −1.5) revealed distinct patterns of EYFP and mCherry in the amygdala. While the CGRP^SPFp^ synaptic terminals were found in the AStr, LA, and medial amygdala (MEA), the CGRP^PBel^ terminals were most abundant in the CeA, and basomedial amygdala (BMA) (Figure 1B). CGRP^SPFp^ neurons did not project to the somatosensory cortex, instead they projected to the auditory cortex and the dorsal regions of the posterior insular cortex (pIC), whereas CGRP^PBel^ neurons projected to the bed nuclei of the stria terminalis (BNST), ventral posteromedial nucleus of the thalamus parvicellular part (VPMpc), parasubthalamic nucleus (PSTN), and the ventral portion of the pIC (Figure S2B).

**Figure 1.**
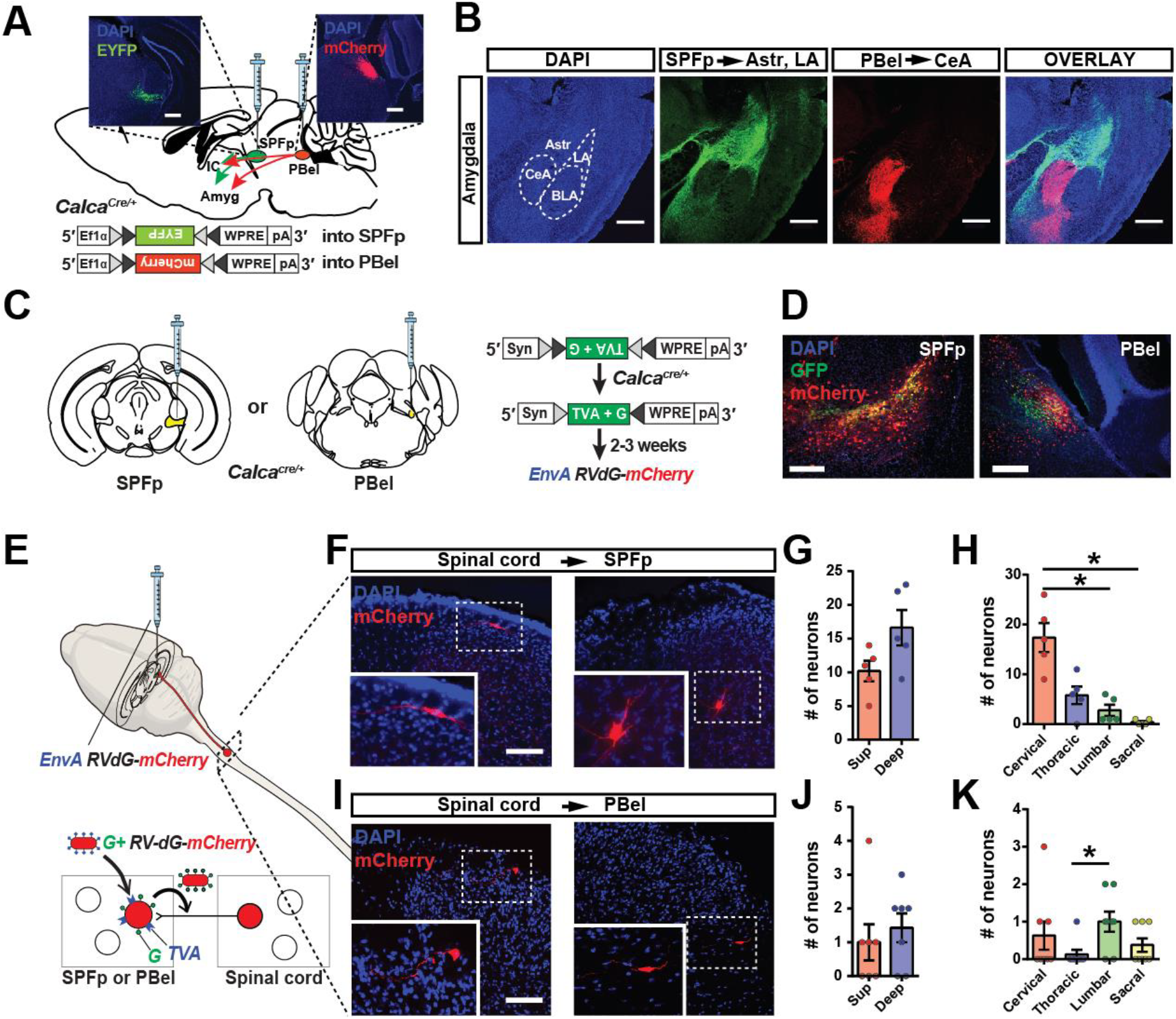
CGRP^SPFp^ and CGRP^PBel^ neurons form spino-thalamo-amygdaloid and spino-parabrachio-amygdaloid pathways. (**A**) Schematic and representative images of Cre-dependent expression of EYFP in the SPFp and mCherry in the PBel of a *Calca^Cre^* mouse. Scale bars, 200 μm. **(B)** The axonal projections from the CGRP^SPFp^ and CGRP^PBel^ neurons are mutually exclusive in the amygdala sub-regions. Scale bars, 500 μm. (**C**) Schematic diagrams and images of Cre-dependent expression of TVA and G in SPFp or PBel neurons of *Calca^Cre^* mice for the rabies tracing. (**D**) Representative images of the SPFp and PBel five days after EnvA-RVdG-mCherry injection. Yellow indicates the starter cells. Scale bars, 200 μm. (**E**) A schematic diagram of identifying presynaptic neurons by monosynaptic rabies tracing. (**F**) Representative images of superficial and deep layer dorsal horn neurons that project to CGRP^SPFp^ neurons.(**G**) The number of spinal dorsal horn neurons in the superficial (Sup) and deep (deep) layers project to the CGRP^SPFp^ neurons. (**H**) The number of spinal cord neurons in different spinal segments that project to the CGRP^SPFp^ neurons.(**I**) Representative images of the superficial dorsal horn and lateral spinal nucleus neurons project to CGRP^PBel^ neurons. (**J**) The number of spinal dorsal horn neurons in the superficial (Sup) and deep (deep) layers project to the CGRP^PBel^ neurons. (**K**) The number of spinal cord neurons in different spinal segments that project to the CGRP^PBel^ neurons. **Statistics** (G) Superficial: 10.20 ± 1.53, Deep: 16.60 ± 2.62 (n = 5). Paired t-test (two-tailed), p = 0.0723. (H) Cervical: 17.40 ± 2.91, Thoracic: 5.80 ± 2.91, Lumbar: 2.80 ± 1.11, Sacral: 0.40 ± 0.24 (n = 5). Repeated measure one-way ANOVA, p =0.0017. Cervical group was significantly different with lumbar (p < 0.05) and sacral (p < 0.05, Tukey’s multiple comparisons). (J) Superficial: 1.00 ± 0.53, Deep: 1.43 ± 0.43 (n = 8). Paired t-test (two-tailed), p = 0.6286. (K) Cervical: 0.63 ± 0.38, Thoracic: 0.13 ± 0.13, Lumbar: 1.00 ± 0.27, Sacral: 0.38 ± 0.18 (n = 8). Repeated measure one-way ANOVA, p =0.1615. Thoracic group was significantly different with lumbar (p < 0.05, Tukey’s multiple comparisons).

To identify upstream brain regions that directly project their axons to the CGRP^SPFp^, or CGRP^PBel^ neurons, we performed cell-type-specific monosynaptic retrograde tracing using pseudotyped rabies virus (Kim et al., 2016). We injected AAV8-hSyn-FLEX-TVA-P2A-GFP-2A-oG into the SPFp or PBel of *Calca^Cre^* mice and then waited three weeks before injecting EnvA-ΔG-rabies-mCherry into the same region (Figure 1C). Five days after the rabies injection, mice were sacrificed to detect starter cells in both regions (Figure 1D) and mCherry-labeled direct upstream neurons in the brain and spinal cord (Figure 1E). Histological analyses revealed that CGRP^SPFp^ neurons received inputs from superficial and deep layers of the spinal dorsal horn in multiple spinal cord segments, but most abundantly from the cervical segment (Figures 1F-1H). By contrast, fewer neurons in the spinal cord project to the CGRP^PBel^ neurons compared to those projecting to the CGRP^SPFp^ neurons (Figures 1I-1K). Other than the spinal cord, CGRP^SPFp^ neurons received inputs from sensory relay areas including the SC, inferior colliculus (IC), vestibular nucleus (VN), and trigeminal spinal nucleus (SpV), as well as other regions such as the hypothalamus and cortex (Figures S3A, S3C, and S3E). CGRP^PBel^ neurons also received projections from sensory relay areas, including the SC, IC, VN, SpV, and the nucleus tractus solitarius (NTS), as well as other areas such as the amygdala (in particular the CeA) and the hypothalamus including the lateral hypothalamus (LHA), zona inserta (ZI), and PSTN (Figures S3B, S3D, and S3F). Thus, these two populations of CGRP neurons may receive multimodal sensory inputs.

Our results show that both CGRP^SPFp^ and CGRP^PBel^ neurons receive monosynaptic inputs from the spinal dorsal horn, other sensory-related regions, hypothalamus, and amygdala (CGRP^PBel^ neurons in particular). However, in terms of output patterns, CGRP^SPFp^ neurons project to the LA and AStr, while CGRP^PBel^ neurons project to the CeA, thereby forming complementary parallel sensory pathways to the amygdala.

### CGRP^SPF^p and CGRP^PBel^ neurons are activated by multimodal nociceptive stimuli

Next, we investigated the response of CGRP^SPFp^ and CGRP^PBel^ neurons to multimodal nociceptive stimuli, such as mechanical, thermal, and inflammatory stimuli using the fiber photometry *in vivo* calcium imaging (Figure 2A). AAV-DIO-GCaMP6m was injected into the SPFp or PBel of *Calca^Cre^* mice, and an optic fiber (400 μm, 0.37 NA) was implanted above the injection site (Figures 2B and 2C). Previous *in vivo* electrophysiology studies have shown that nociceptive signals can be conveyed from the spinal cord to the brain under anesthesia (Gauriau and Bernard, 2004b; Peschanski et al., 1981). Therefore, we performed the fiber photometry experiments under light anesthesia to remove other emotional and movement artifacts.

**Figure 2.**
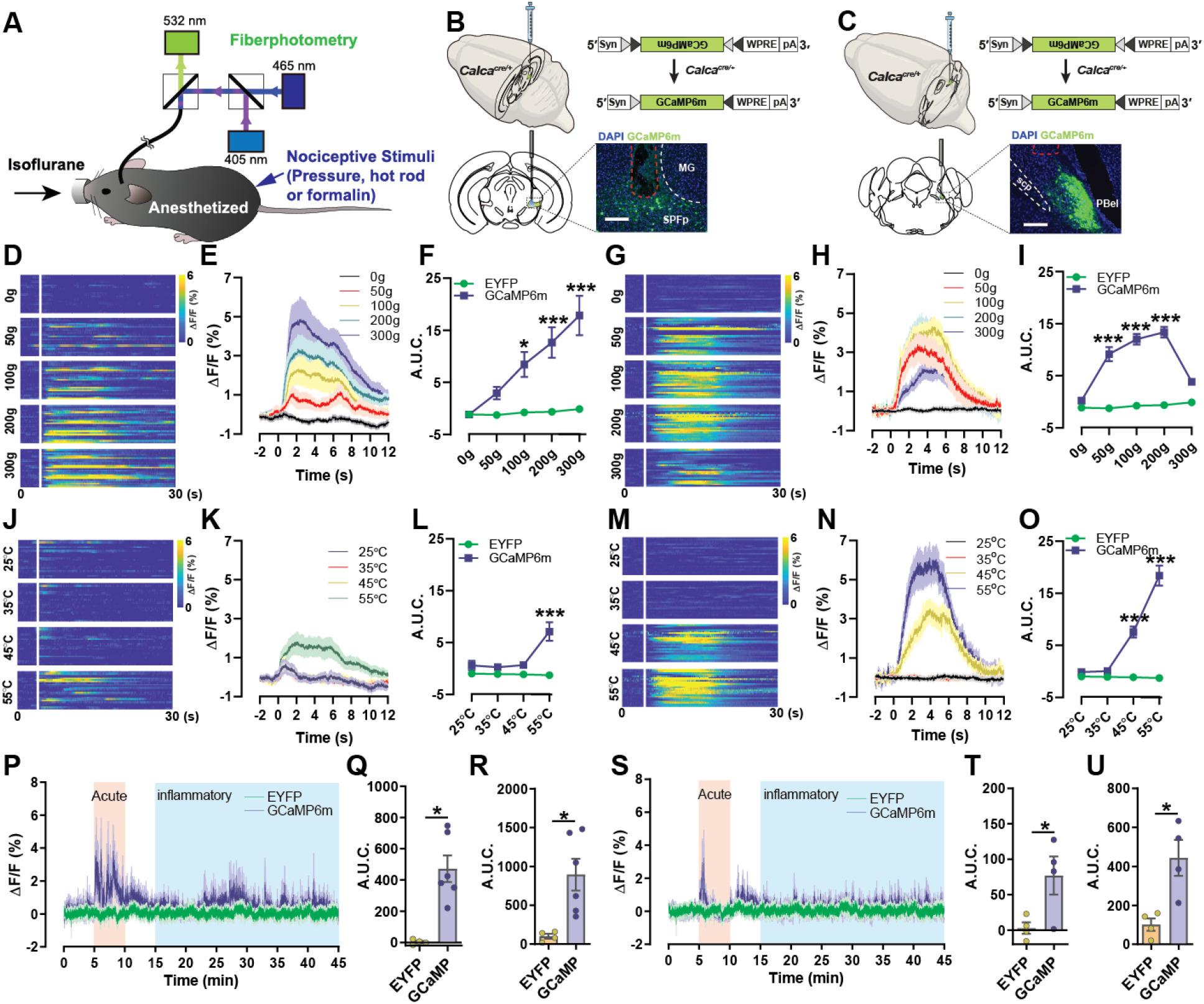
CGRP^SPFp^ and CGRP^PBel^ neurons are activated by multimodal nociceptive stimuli. (**A**) A diagram of the fiber photometry calcium imaging experiment in an anesthetized mouse. (**B and C**) A schematic and representative image of the Cre-dependent expression of GCaMP6m in the *Calca^Cre^* mice with an optical fiber implanted (red dashed line in the overlay image indicates fiber track) above the SPFp and PBel. Scale bars, 200 μm. (**D-F**) Intensity-dependent calcium activity increase in CGRP^SPFp^ neurons in response to a pressure meter (0, 50, 100, 200, and 300 g). (**G-I**) Intensity-dependent calcium activity increase in CGRP^PBel^ neurons to a pressure meter. (**J-L**) Intensity-dependent calcium increase in CGRP^SPFp^ neurons in response to a temperature-controlled rod (25, 35, 45, or 55 °C). (**M-O**) Intensity-dependent calcium activity increase in CGRP^PBel^ neurons in response to a temperature-controlled rod. (**P**) Calcium signal increases in CGRP^SPFp^ neurons following subcutaneous injection of formalin (4%) into the paw. (**Q**) Area under curve (A.U.C.) quantification of the CGRP^SPFp^ neuronal activity during the acute phase (5– 10 min) of formalin response. (**R**) A.U.C. quantification of the CGRP^SPFp^ neuronal activity during the inflammatory phase (15–45 min) of formalin injection. (**S**) Calcium signal increases in CGRP^PBel^ neurons in response to subcutaneous injection of formalin (4%) into the paw. (**T**) A.U.C. quantification of the CGRP^PBel^ neuronal activity during the acute phase (5–10 min) of formalin response. (**U**) A.U.C. quantification of the CGRP^PBel^ neuronal activity during the inflammatory phase (15–45 min) of formalin response. **Statistics** (F) Repeated measure two-way ANOVA showed significance in interaction (F (4, 152) = 9.838, p < 0.0001), intensity (F (4, 152) = 12.28, p < 0.0001) and group (F (1, 38) = 16.01, p = 0.0003). 100 (p < 0.05), 200 (p < 0.0001) and 300 g (p < 0.0001) points were significantly different between EYFP and GCaMP6m with Sidak’s multiple comparisons test. (I) Repeated measure two-way ANOVA showed significance in interaction (F (4, 248) = 15.08, p < 0.0001), intensity (F (4, 248) = 15.17, p < 0.0001) and group (F (1, 62) = 64.40, p < 0.0001). 50 (p < 0.0001), 100 (p < 0.0001) and 200 g (p < 0.0001) points were significantly different between EYFP and GCaMP6m with Sidak’s multiple comparisons test. (L) Repeated measure two-way ANOVA showed significance in interaction (F (3, 114) = 10.97, p <0.0001), intensity (F (3, 114) = 9.61, p < 0.0001) and group (F (1, 38) = 11.12, p = 0.0019). 55°C (p < 0.0001) was significantly different between EYFP and GCaMP6m with Sidak’s multiple comparisons test. (M) Repeated measure two-way ANOVA showed significance in interaction (F (3, 186) = 23.96, p < 0.0001), intensity (F (3, 186) = 22.67, p < 0.0001) and group (F (1, 62) = 46.05, p < 0.0001). 45 and 55°C (both p < 0.0001) points were significantly different between EYFP and GCaMP6m with Sidak’s multiple comparisons test. (Q) EYFP: 2.86 ± 8.00 (n = 4 mice), GCaMP6m: 471.9 ± 85.53 (n = 6 mice). Unpaired t-test (two-tailed), p = 0.0024. (R) EYFP: 100.60 ± 31.89 (n = 4), GCaMP6m: 894.80 ± 203.40 (n = 6). Unpaired t-test (two-tailed), p = 0.0145. (K) EYFP: 6.18 ± 4.05 (n = 4), GCaMP6m: 77.00 ± 26.83 (n = 4). Unpaired t-test (two-tailed), p = 0.0381. (L) EYFP: 100.60 ± 31.89 (n = 4), GCaMP6m: 443.50 ± 92.46 (n = 4). Unpaired t-test (two-tailed), p = 0.0127.

Various intensities of mechanical stimuli (0, 50, 100, 200, and 300 g of pressure delivered by a pressure meter, not von Frey hairs) were applied for 5 seconds to the ipsi-or contra-lateral paws or tail, resulting in intensity-dependent increases in calcium signals (Area under curve analysis; A.U.C.) in both CGRP^SPFp^ (Figures 2D-2F) and CGRP^PBel^ neurons (Figures 2G-2I), but there was a significantly decreased calcium response in the CGRP^PBel^ neurons by 300 g stimulation which created the inverted U-shaped intensity response curve. Nevertheless, CGRP^SPFp^ and CGRP^PBel^ neurons display different dynamics to noxious stimuli. CGRP^PBel^ neurons reached maximum response at lower stimulation intensity compared to the CGRP^SPFp^ neurons (Figures S4A and S4B), but the latter responded faster to the stimuli, which was observed by greater initial rise slope of calcium peak in the CGRP^SPFp^ neurons compared to the CGRP^PBel^ neurons (Figure S4C). Although contralateral stimulation evoked greater responses than the ipsilateral stimulation, both the CGRP^SPFp^ and CGRP^PBel^ neurons were robustly activated by noxious stimuli from both sides, contrary to the conventional lateralized ascending sensory pain pathways. (Figures S4D-S4G).

Heat stimuli (25, 35, 45, and 55°C) also induced intensity-dependent calcium increases in both CGRP^SPFp^ (Figure 2J-2L) and CGRP^PBel^ neurons (Figure 2M-2O). The maximum calcium peak values were higher overall in CGRP^PBel^ neurons than in CGRP^SPFp^ neurons, both for contralateral (Figure S4H) and ipsilateral stimulations (Figure S4I). However, the CGRP^SPFp^ neurons again responded faster to 55 °C stimuli than CGRP^PBel^ neurons (Figure S4J). Although the difference between ipsilateral and contralateral stimulation was observed in CGRP^SPFp^ neurons, these neurons were robustly activated by thermal noxious stimuli from both sides (Figures S4K–S4N). Interestingly, calcium responses induced by mechanical stimuli were much stronger than those induced by thermal stimuli in CGRP^SPFp^ neurons; the opposite was observed for CGRP^PBel^ neurons. These results indicate that the CGRP^SPFp^ and CGRP^PBel^ neurons may play different roles in conveying mechanical and thermal pain.

To assess the effects of inflammatory pain, we injected 10 μl of 4% formalin into the contralateral forepaw, and calcium activity was recorded with fiber photometry under light anesthesia (Figures 2P-2U). We observed increases in activity during both the initial acute pain phase (5–10 min; Figures 2Q and 2T) and the inflammatory phase (15–45 min; Figures 2R and 2U). Activation of these neurons by formalin was confirmed by Fos immunostaining, which again showed that bilateral CGRP^SPFp^ and CGRP^PBel^ neurons were activated by unilateral formalin injection with slightly higher response in the contralateral neurons (Figures S5A-S5G).

Our data indicate that multimodal nociceptive stimuli bilaterally activate CGRP^SPFp^ and CGRP^PBel^ neurons and that these two populations differentially respond to nociceptive inputs of distinct modalities. Moreover, there is a potential feedback inhibitory circuit in the CGRP^PBel^ neurons, as we observed activity decreases in response to 300 g of mechanical pain and during the acute phase of the formalin test.

### Pain-induced synaptic plasticity change in CGRP^SPFp^ and CGRP^PBel^ neurons

As we observed that noxious stimuli activate CGRP^SPFp^ and CGRP^PBel^ neurons, we hypothesized that pain might alter the glutamatergic synaptic strength of these neurons. We performed *ex vivo* electrophysiology to test pain-induced synaptic plasticity changes. To induce pain, we injected 50 μl of 5% formalin into the upper lip. Twenty-four hours after the injection, we prepared acute brain slices that contained the SPFp (Figure 3A) or PBel (Figure 3I). We then measured the AMPA/NMDA ratio, an index of glutamatergic synaptic strength, for both CGRP^SPFp^ and CGRP^PBel^ neurons using the whole-cell patch-clamp recording. For both CGRP^SPFp^ (Figures 3B and 3C) and CGRP^PBel^ neurons (Figures 3J and 3K), the AMPA/NMDA ratio increased (indicating long-term potentiation) in mice treated with formalin compared to controls. No differences were observed in the paired-pulse ratio (which indicates a presynaptic mechanism) for CGRP^SPFp^ neurons (Figures 3D and 3E), but an increase in this ratio was found for CGRP^PBel^ neurons (Figures 3L and 3M). To determine whether the AMPA current was affected, we recorded AMPA-mediated mini EPSCs (Figures 3F and 3N). mEPSC amplitude (Figure 3G), but not frequency (Figure 3H), increased in CGRP^SPFp^ neurons of mice subjected to pain compared with controls. For CGRP^PBel^ neurons, the pain did not affect mEPSC amplitude (Figure 3O) but decreased frequency (Figure 3P).

**Figure 3.**
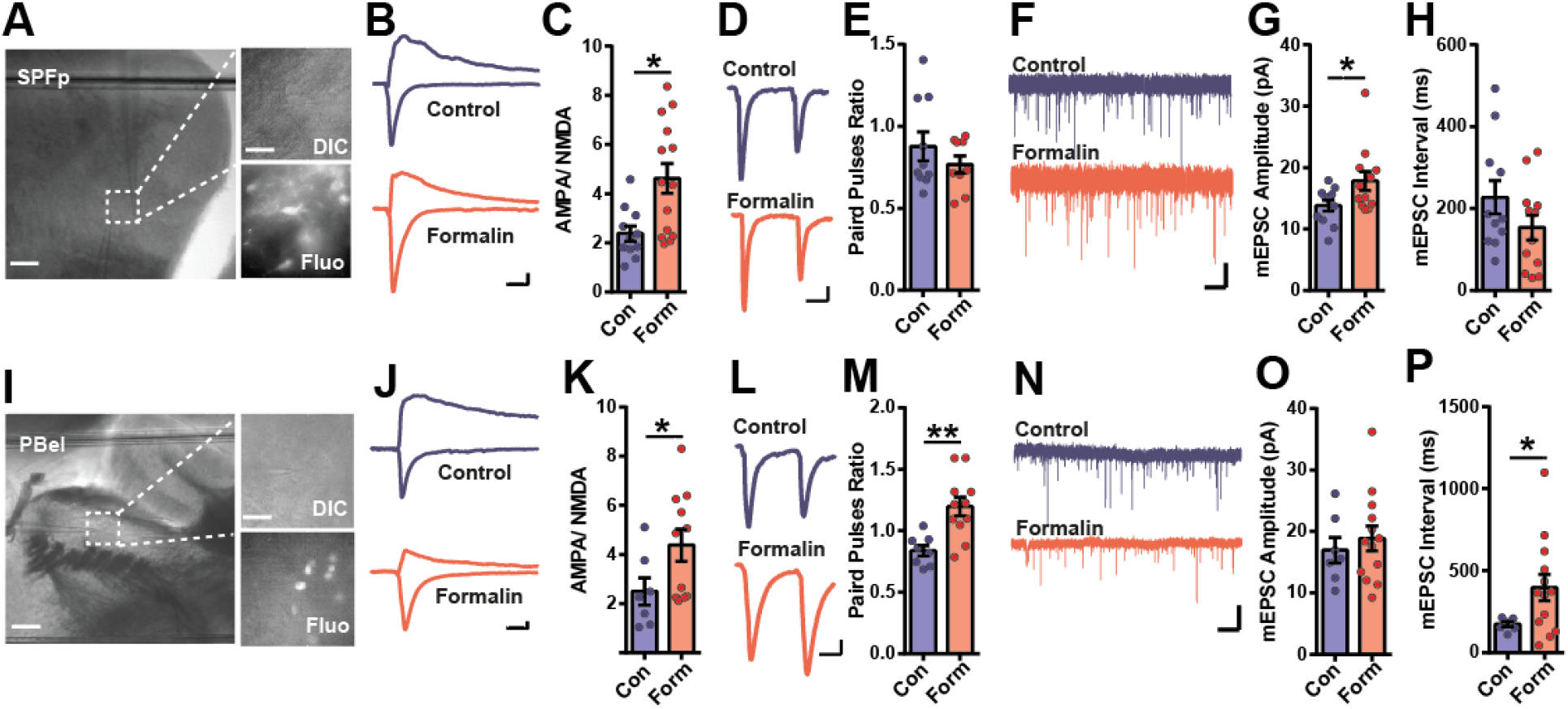
Pain-induced synaptic plasticity changes in CGRP^SPFp^ and CGRP^PBel^ neurons. (**A**) A representative image of a brain slice including SPFp used for the whole-cell patch-clamp experiment. Enlarged images show the fluorescence from Cre:GFP-expressing CGRP neurons. Scale bars, 100 μm in left images, and 10 μm in enlarged images. (**B**) Example traces of AMPA and NMDA EPSCs in control (blue) and formalin (red) injected groups. Scale bars, 20 ms, and 50 pA. (**C**) The AMPA/NMDA ratio was increased in the formalin injected group compared with the control group. (**D and E**) No significant differences in the paired-pulse ratio were observed between the formalin-injected group and the control group. Scale bars, 20 ms, and 50 pA. (**F**) Example traces of mEPSCs in the control (blue) and formalin (red) injected group. Scale bars, 1 s, and 20 pA. (**G**) mEPSC amplitude was increased in the formalin injected group compared with the control group. (**H**) mEPSC interval was not changed in the formalin injected group compared with the control group. (**I**) A representative image of a brain slice including PBel for the whole-cell patch-clamp experiment. Enlarged images are the PBel cells with and without fluorescence. Cells with fluorescence are CGRP neurons. Scale bars, 100 μm in left images, and 10 μm in enlarged images. (**J**) Example traces of AMPA and NMDA EPSCs in control (blue) and formalin (red) injected groups. Scale bars, 20 ms, and 20 pA. (**K**) The AMPA/NMDA ratio was increased in the formalin injected group compared with the control group. (**L and M**) The formalin injected group display an increased paired-pulse ratio compared to the control group. Scale bars, 20 ms, and 50 pA. (**N**) Example traces of mEPSCs in the control (blue) and formalin (red) injected group. Scale bars. 1 s and 50 pA. (**O**) mEPSC amplitude was not changed in the formalin injected group compared with the control group. (**P**) mEPSC interval increased in the formalin injected group compared with the control group. **Statistics** (C) SPFp; Control: 2.38 ± 0.31 (n = 11 cells), formalin: 4.62 ± 0.61 (n = 14 cells). Unpaired t test (two-tailed), p = 0.0063. (E) SPFp; Control: 0.88 ± 0.09 (n = 10 cells), formalin: 0.77 ± 0.05 (n = 9 cells). Unpaired t test (two-tailed), p = 0.2957. (G) SPFp; Control: 13.81 ± 0.90 (n = 11 cells), formalin: 17.83 ± 1.56 (n = 12 cells). Unpaired t test (two-tailed), p = 0.0390. (H) SPFp; Control: 227.4 ± 40.90 (n = 11 cells), formalin: 154.2 ± 30.80 (n = 12 cells). Unpaired t test (two-tailed), p = 0.1687. (K) PBel; Control: 2.49 ± 0.56 (n = 7 cells), formalin: 4.38 ± 0.65 (n = 11 cells). Unpaired t test (two-tailed), p = 0.0437. (M) PBel; Control: 0.84 ± 0.04 (n = 8 cells), formalin: 1.20 ± 0.77 (n = 11 cells). Unpaired t test (two-tailed), p = 0.001. (O) PBel; Control: 16.93 ± 2.08 (n = 7 cells), formalin: 18.85 ± 2.04 (n = 13 cells). Unpaired t test (two-tailed), p = 0.5196. (P) PBel; Control: 173.2 ± 13.89 (n = 7 cells), formalin: 396.5 ± 80.66 (n = 13 cells). Unpaired t test (two-tailed), p = 0.0175.

These data together suggest that CGRP^SPFp^ and CGRP^PBel^ neuronal synapses differentially increase the strength of glutamatergic signaling in response to pain.

### The transcriptome profiling of CGRP^SPFp^ and CGRP^PBel^ neurons

To further investigate whether CGRP^SPFp^ and CGRP^PBel^ neurons exhibit specific transcriptomic profiles associated with the affective pain perception, we conducted cell-type-specific transcriptomic profiling of CGRP neurons in the SPFp and PBel regions. The *Calca^CreER^* mouse line was crossed with the RiboTag mouse line (Sanz et al., 2009) that has a floxed allele of hemagglutinin (HA)-tagged *Rpl22* gene. As a result, the HA-tagged ribosomal protein, RPL22 is Cre-dependently expressed in the CGRP-expressing neurons. After fresh brain tissues containing the SPFp or PBel region were collected and homogenized, the ribosome-associated transcriptome was captured by immunoprecipitation with anti-HA antibody-conjugated magnetic beads. Precipitated / unprecipitated total RNAs were sequenced to profile active transcriptome enriched / deenriched in CGRP^SPFp^ and CGRP^PBel^ neurons. RNA sequencing results revealed that the *Calca* gene that encodes CGRP is highly enriched in both CGRP^SPFp^ and CGRP^PBel^-specific transcriptome, which served as a positive control. Moreover, genes encoding neuropeptide were specifically enriched in CGRP^PBel^ neurons (Figure 4D), and genes encoding markers for inhibitory neurons or glia were de-enriched in both regions (Figure 4E), confirming that these neurons are glutamatergic neurons. Notably, several pain-related genes, in particular those encoding membrane proteins, were enriched in both populations (Figures 4A-4C). Interestingly, genes associated with affective pain disorders, such as *Scn9a*, and *Faah* for congenital insensitivity to pain (CIP), and *Cacna1a* for migraine (Nassar et al., 2004; van den Maagdenberg et al., 2004; Cravatt and Lichtman, 2004) are highly enriched in these neurons (Figures 4A-4C). Expression of the proteins encoded by these genes (Na_v_1.7, Ca_v_2.1, and FAAH) in the SPFp and PBel was confirmed via immunohistochemistry (Figures 4F-4H). These data indicate that both CGRP^SPFp^ and CGRP^PBel^ neurons express genes involved in pain perception, further supporting our results that these neurons form ascending pain pathways.

**Figure 4.**
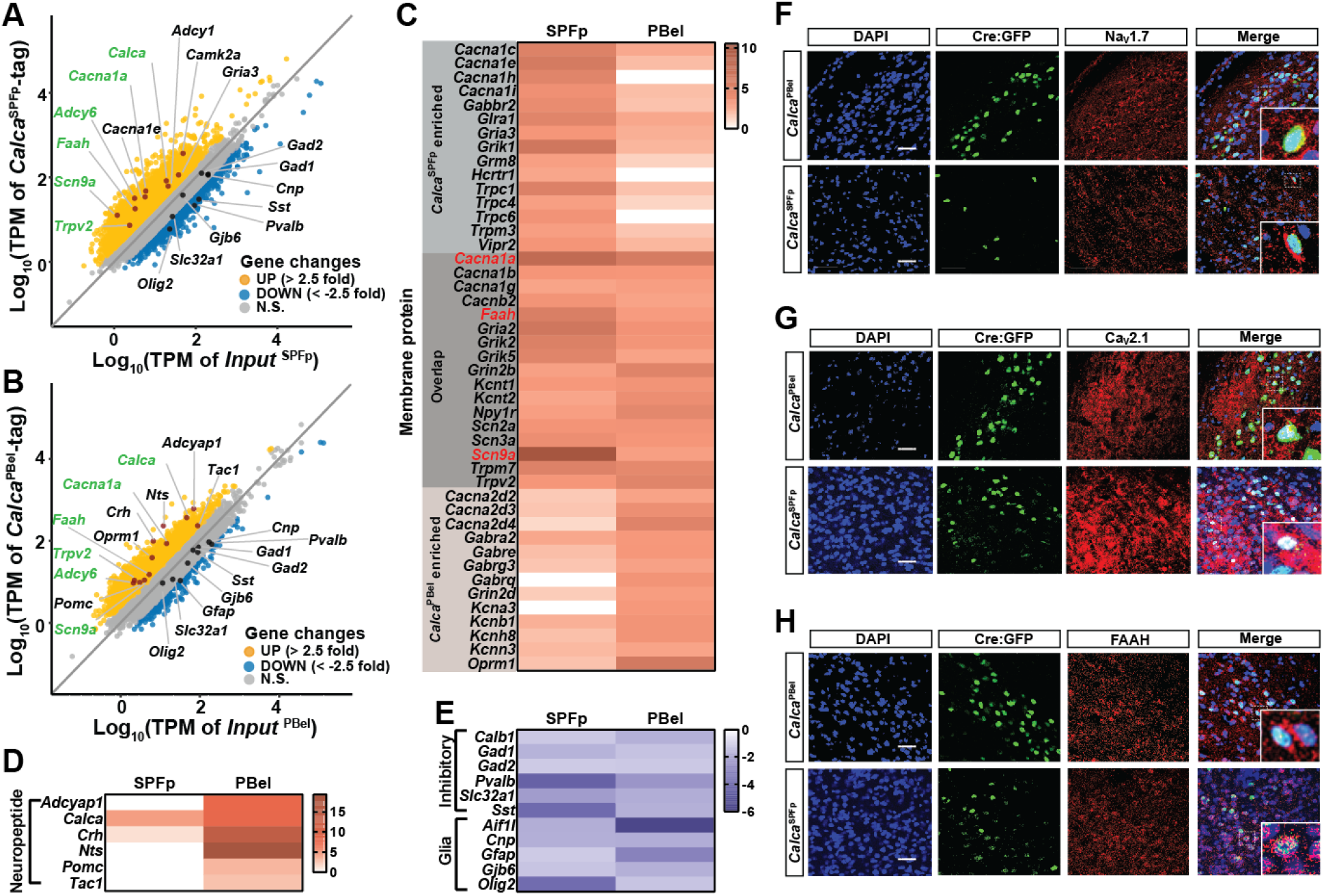
Active transcriptome profiling of CGRP^SPFp^ and CGRP^PBel^ neurons. (**A**) Correlation plot showing expression (transcript per million, TPM in log 10 scale) of genes enriched in CGRP^SPFp^ neurons compared with the total SPFp inputs using RiboTag transcriptome profiling. Up and down-regulated genes were separated based on 2.5 or −2.5-fold enrichment. (**B**) Correlation plot of the transcriptome profiles of CGRP^PBel^ neurons versus total PBel inputs. (**C**) Heatmaps showing fold changes of genes in the SPFp and PBel that encode membrane proteins. (**D**) Heatmaps showing fold changes of genes in the SPFp and PBel that encode neuropeptides, (**E**) Heatmaps showing fold changes of genes in the SPFp and PBel that encode markers of inhibitory neuron or glia. (**F-H**) Co-expression of NaV1.7 (**F**), CaV2.1 (**G**), or FAAH (**H**) with CGRP by double IHC. Scale bars, 50 μm.

### CGRP^SPF^p and CGRP^PBel^ neurons are activated by multisensory innate threat stimuli

Our retrograde tracing results indicate that both CGRP^SPFp^ and CGRP^PBel^ neurons receive inputs from areas conveying sensory information from multiple sensory modalities, such as visual stimulus via the SC, auditory stimulus via the IC, gustatory stimulus via NTS, and nociceptive stimuli via the SC and SpV. We, therefore, examined whether they are also activated by multisensory innate threat stimuli. We used the fiber photometry system to measure neural activity in response to five different aversive sensory stimuli. AAV-DIO-GCaMP7s was injected into the SPFp or PBel of *Calca^Cre^* mice, and an optic fiber (400 μm, 0.37 NA) was implanted above the injection site to measure calcium activity of CGRP^SPFp^ or CGRP^PBel^ neurons (Figures 5A and 5B). A somatosensory stimulus was first tested by applying a foot shock in a cued fear conditioning test (Figure 5C). We associated a non-aversive low-volume tone (70 dB) with the shock to minimize the tone’s aversive effect (Figure S6A). For both CGRP^SPFp^ and CGRP^PBel^ neurons, immediate increases in neural activity were detected in response to the 2-s foot shock, but not during habituation or during the cue test (when the tone was on; Figures 5D–5E). Freezing was observed during the cue test, indicating that fear memory was formed (Figures S6B and S6C). To assess the auditory threat, an intense sound (85 dB) was delivered for 2 s (Figure 5F). Time-locked calcium responses were detected at the onset of an 85-dB intense sound, but not a 70-dB sound for both CGRP^SPFp^ (Figure 5G) and CGRP^PBel^ neurons (Figure 5H). For an innate visual stimulus, a 2-s looming stimulus was given three times with 10-s intervals. Both CGRP^SPFp^ (Figure 5J) and CGRP^PBel^ neurons (Figure 5K) displayed an increase of activity in response to the looming (large disk) compared to the control (small disk) stimulus. As an innate olfactory stimulus test, we exposed mice to a cotton swab soaked with trimethylthiazoline (TMT; Figure 5L). CGRP^SPFp^ neurons did not respond to TMT (Figure 5M), whereas CGRP^PBel^ neurons exhibited a slight increase in activity (Figure 5N). Finally, a gustatory stimulus was administered by exposing mice to quinine (vs. water; Figure 5O). When overnight water restricted mice licked quinine solution (0.5 mM), CGRP^SPFp^ neurons did not respond, compared with water controls (Figure 5P), but CGRP^PBel^ neurons exhibited an increase in calcium activity (Figure 5Q). The calcium peak amplitude analysis shows that the calcium responses between the CGRP^SPFp^ and CGRP^PBel^ neurons were not significantly different by somatosensory and auditory stimuli (Figures S7A and S7B). However, the CGRP^SPFp^ neurons showed a greater response to the visual stimulus compared to the CGRP^PBel^ neurons (Figure S7C). In contrast, the CGRP^PBel^ neurons showed greater responses to the olfactory and gustatory stimuli compared to the CGRP^SPFp^ neurons (Figures S7D and S7E).

**Figure 5.**
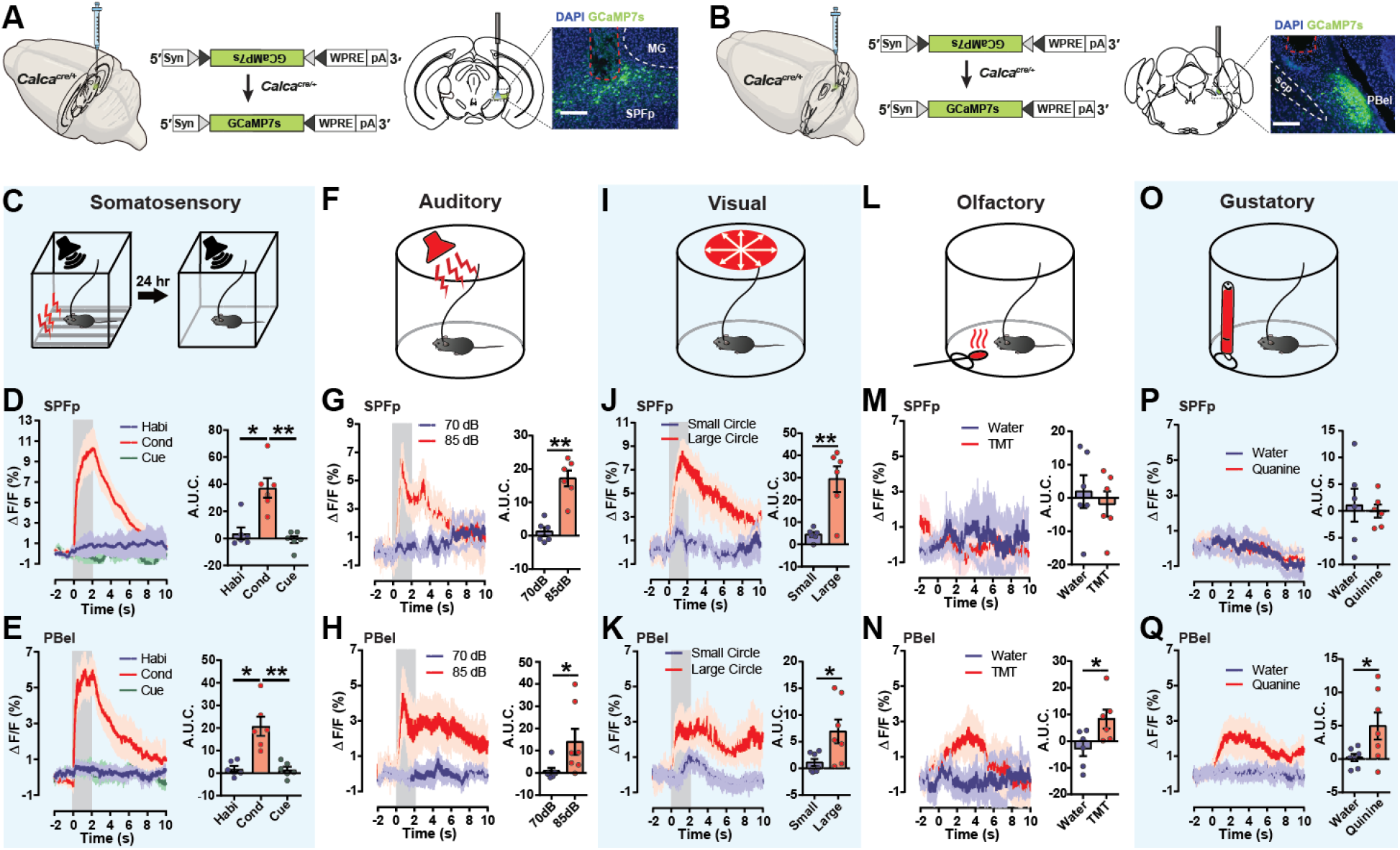
CGRP^SPFp^ and CGRP^PBel^ neurons are activated by multimodal sensory threat stimuli. (**A and B**) A schematic and representative images of Cre-dependent expression of GCaMP7s in the CGRP^SPFp^ (**A**) and CGRP^PBel^ (**B**) neurons for fiber photometry. Scale bars, 200 μm. (**C**) Cued fear conditioning test with low volume tone (72 dB) was performed to examine the responses of CGPR neurons to the somatosensory aversive stimulus (2-s, 0.6 mA electric foot shock). (**D and E**) CGRP^SPFp^ (**D**) and CGRP^PBel^ (**E**) neurons were activated by the electric foot shock during the conditioning period, but not habituation, nor the cued retrieval period. Left, averaged calcium traces. Right, A.U.C. quantification. (**F**) Intense sound (85 dB) was used as an aversive auditory stimulus, with a70-dB low-intensity sound as a control. (**G and H**) CGRP^SPFp^ (**G**) and CGRP^PBel^ (**H**) neurons were activated by the intense sound. Left, averaged calcium traces. Right, A.U.C. quantification. (**I**) A large looming disk was used as an aversive visual stimulus, with a small disk as a control. (**J and K**) CGRP^SPFp^ (**J**) and CGRP^PBel^ (**K**) neurons were activated by the large looming disk. Left, averaged calcium traces. Right, A.U. C. quantification. (**L**) TMT-soaked cotton was used as an aversive olfactory stimulus, with water as a control. (**M**) There was no activity change in CGRP^SPFp^ neurons when the animal approached the TMT-soaked cotton. Left, averaged calcium traces. Right, A.U. C. quantification. (**N**) CGRP^PBel^ neurons were activated when the animal approached the TMT-soaked cotton. Left, averaged calcium traces. Right, A.U. C. quantification. (**O**) Quinine solution (0.5 mM) was introduced as an aversive gustatory stimulus (water was the control) (**P**) Quinine did not affect CGRP^SPFp^ neurons. Left, averaged calcium traces. Right, A.U. C. quantification. (**Q**) CGRP^PBel^ neurons were activated by quinine solution (0.5 mM). Left, averaged calcium traces. Right, A.U.C. quantification. **Statistics** (D) SPFp; Habituation: 3.63 ± 4.45, conditioning: 37.31 ± 7.18, cue test: −0.92 ± 2.57 (n = 6). Repeated measure one-way ANOVA, p = 0.0029. Conditioning was significantly different with habituation (p < 0.05) and cue test (p <0.01, Tukey’s multiple comparisons). (E) PBel; SPFp; Habituation: 1.85 ± 1.33, conditioning: 20.73 ± 4.25, cue test: 1.49 ± 1.35 (n = 6). Repeated measure one-way ANOVA, p = 0.0022. Conditioning was significantly different with habituation (p < 0.05) and cue test (p <0.01, Tukey’s multiple comparisons). (G) SPFp; 70 dB: 1.226 ± 1.27, 85 dB: 17.14 ± 2.38 (n = 6). Paired t test (two-tailed), p = 0.0018. (H) PBel; 70 dB: 0.78 ± 1.46, 85 dB: 14.06 ± 5.86 (n = 7). Paired t test (two-tailed), p = 0.0437. (J) SPFp; Small disk: 4.41 ± 1.11, large disk: 29.30 ± 5.81 (n = 6). Paired t test (two-tailed), p = 0.0091. (K) PBel; Small disk: 1.08 ± 0.69, large disk: 6.93 ± 2.23 (n = 7). Paired t test (two-tailed), p = 0.0357. (M) SPFp; Water: 1.93 ± 4.88, TMT: −1.82 ± 3.78 (n = 6). Paired t test (two-tailed), p = 0.4411. (N) PBel; Water: −2.72 ± 2.84, TMT: 8.28 ± 3.57 (n = 6). Paired t test (two-tailed), p = 0.0145. (P) SPFp; Water: 1.10 ± 3.09, quinine: −0.01 ± 1.22 (n = 6). Paired t test (two-tailed), p = 0.7707. (Q) PBel; Water: 0.2209 ± 0.83, quinine: 4.96 ± 2.01 (n = 7). Paired t test (two-tailed), p = 0.0377.

Our results indicate that the CGRP^SPFp^ and CGRP^PBel^ neurons are both involved in the perception of innate multisensory threat but respond differently to inputs from distinct modalities. CGRP^PBel^ neurons were activated by all five aversive sensory stimuli, whereas the CGRP^SPFp^ neurons were only activated by somatosensory, auditory, and visual aversive stimuli.

### Silencing CGRP neurons attenuates responses to multimodal threat stimuli

Our rabies tracing and fiber photometry results imply that CGRP^SPFp^ and CGRP^PBel^ neurons are critically involved in innate threat perception. Thus, we next tested whether these neurons are necessary for innate threat perception. We silenced these neurons by bilateral injection of AAV-DIO-TetTox::GFP into the SPFp or PBel of *Calca^Cre^* mice and measured behavioral responses to pain stimuli, and multimodal aversive threat stimuli (Figure 6A). We first performed the formalin assay to test the affective pain perception (Figure 6B). Following injection of 4% formalin into the forepaw, mice in which CGRP^SPFp^ or CGRP^PBel^ neurons were silenced spent less time licking the injected paw (Figure 6C). In addition, the CGRP^SPFp^ silenced group exhibited decreased thermal sensitivity in the 55 °C hot plate test (Figure S8A and S8B), decreased mechanical sensitivity in the electronic von Frey test (Figures S8C and S8D), and decreased freezing in response to the contextual fear conditioning test (Figures S8E and S8F). Interestingly, previous results with CGRP^PBel^-silenced mice by TetTox exhibited no changes in thermal or mechanical thresholds but decreased freezing in the fear conditioning test (Han et al., 2015), suggesting that both CGRP^SPFp^ and CGRP^PBel^ neurons are necessary for affective pain perception. However, CGRP^SPFp^ neurons, not CGRP^PBel^ neurons, are also necessary for sensory pain perception. The elevated plus maze (EPM) test shows that silencing the CGRP^SPFp^ or CGRP^PBel^ neurons decreased anxiety-like behaviors in mice (Figures S8G and S8H).

**Figure 6.**
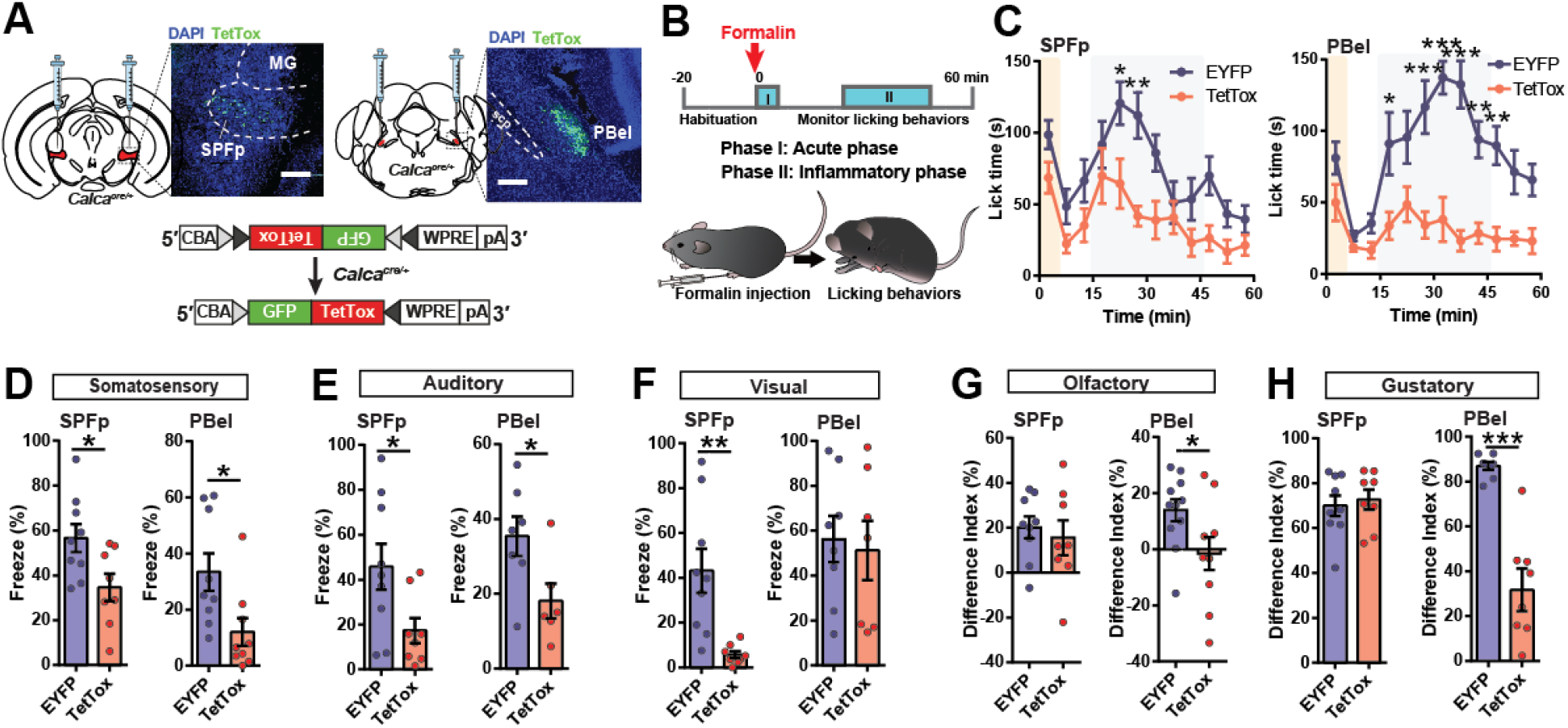
Silencing CGRP^SPFp^ or CGRP^PBel^ neuronal activities attenuates perception of affective pain and multisensory threat stimuli. (**A**) Schematics and representative images of Cre-dependent TetTox expression targeting CGRP^SPFp^ or CGRP^PBel^ neurons. Scale bars, 200 μm. (**B**) A schematic diagram of the Formalin assay to test inflammatory pain responses. (**C**) The CGRP^SPFp^ and CGRP^PBel^ silenced groups displayed significantly alleviated inflammatory pain responses. (**D**) Both the CGRP^SPFp^ and CGRP^PBel^ silenced groups froze less in response to electric foot shock (2-s, 0.6 mA) compared to controls. (**E**) Both the CGRP^SPFp^ and CGRP^PBel^ silenced groups froze less in response to 85-dB intense sound compared with controls. (**F**) In response to a looming aversive visual stimulus, the CGRP^SPFp^ silenced group showed less freezing, whereas there was no difference between EYFP and CGRP^PBel^ silenced groups. (**G**) A two-chamber choice test with TMT- and water-soaked cotton placed at each corner of the chamber was used to test animals’ responses to aversive olfactory stimulus. The CGRP^SPFp^ silenced group, and EYFP controls both avoided the TMT chamber, while the CGRP^PBel^ TetTox group exhibited less aversion to TMT, spending equal amounts of time in the water and TMT chambers. (**H**) A quinine-water choice test was performed to test the animals’ responses to the aversive gustatory stimulus (0.5 mM quinine solution). There was no change in water preference in the CGRP^SPFp^ silenced group, but the CGRP^PBel^ silenced group showed a lower difference index than the control group. **Statistics** (C) SPFp: Repeated measure two-way ANOVA showed no significance in interaction (F (11, 187) = 1.32, p = 0.2148), but in time (F (11, 187) = 9.05, p < 0.0001) and group (F (1, 17) = 9.57, p = 0.0066). 20-25 (p < 0.05) and 25-30 (p < 0.01) min point were significantly different between EYFP and TetTox with Sidak’s multiple comparisons test. PBel: Repeated measure two-way ANOVA showed significance in interaction (F (11, 187) = 3.45, p = 0.0002), time (F (11, 187) = 5.97, p < 0.0001) and group (F (1, 17) = 53.24, p < 0.0001). 15-20 (p < 0.05), 25-30 (p < 0.0001), 30-35 (p < 0.0001), 35-40 (p < 0.0001), 40-45 (p < 0.01), 45-50 (p < 0.01) min point were significantly different between EYFP and TetTox with Sidak’s multiple comparisons test. (D) SPFp; EYFP: 56.62 ± 6.21 % (n = 9), TetTox: 34.65 ± 6.11 % (n = 8). Unpaired t test (two-tailed), p = 0.0240. PBel; EYFP: 33.43 ± 6.66 % (n = 9), TetTox: 12.04 ± 4.88 % (n = 9). Unpaired t test (two-tailed), p = 0.0197. (E) SPFp; EYFP: 45.74 ± 10.26 % (n = 9), TetTox: 17.32 ± 5.58 % (n = 8). Unpaired t test (two-tailed), p = 0.0332. PBel; EYFP: 35.39 ± 5.29 % (n = 7), TetTox: 18.14 ± 4.66 % (n = 6). Unpaired t test (two-tailed), p = 0.0350. (F) SPFp; EYFP: 43.12 ± 9.82 % (n = 9), TetTox: 5.73 ± 1.57 % (n = 8). Unpaired t test (two-tailed), p = 0.0029. PBel; EYFP: 56.20 ± 10.33 % (n = 8), TetTox: 51.22 ± 13.21 % (n = 7). Unpaired t test (two-tailed), p = 0.0029. (G) SPFp; EYFP: 20.12 ± 4.91 % (n = 9), TetTox: 15.55 ± 7.75 % (n = 8). Unpaired t test (two-tailed), p = 0.6175. PBel; EYFP: 13.87 ± 3.94 % (n = 11), TetTox: −1.53 ± 5.89 % (n = 10). Unpaired t test (two-tailed), p = 0.0396. (H) SPFp; EYFP: 69.96 ± 4.64 % (n = 9), TetTox: 72.66 ± 4.48 % (n = 8). Unpaired t test (two-tailed), p = 0.6823. PBel; EYFP: 86.87 ± 1.77 % (n = 8), TetTox: 31.66 ± 9.45 % (n = 7). Unpaired t test (two-tailed), p < 0.0001.

To test the role of these neurons in multisensory threat perception, the same group of mice were subjected to multiple aversive sensory threat cues, as described in Figure 5. Levels of immediate freezing in response to the aversive somatosensory stimulus (2-s, 0.6 mA electric foot shock) were significantly reduced in both the CGRP^SPFp^ and CGRP^PBel^ TetTox groups compared to the EYFP control groups (Figure 6D). In the auditory threat test with 85-dB intense sound, EYFP control mice displayed freezing behavior, but freezing levels were reduced in both the CGRP^SPFp^ and CGRP^PBel^ TetTox groups (Figure 6E). Defensive behaviors (freezing) were also attenuated in response to a looming visual stimulus in the CGRP^SPFp^ TetTox group compared with controls, but no difference in freezing was observed between the CGRP^PBel^ TetTox group and controls (Figure 6F). Interestingly, CGRP^PBel^ neurons were activated by looming (Figure 5K), but their silencing was not enough to attenuate the animals’ response to a visual threat, indicating that they play a less significant role in transmitting aversive visual stimulus to the amygdala compared to the CGRP^SPFp^ neurons. The aversive olfactory test was performed using a two-chamber system, with one chamber containing water-soaked cotton and the other containing TMT-soaked cotton. Silencing the CGRP^SPFp^ neurons did not affect the perception of the aversive olfactory cue, as these mice and EYFP controls both avoided the TMT chamber, while the CGRP^PBel^ TetTox group exhibited no aversion to TMT, spending equal amounts of time in the water and TMT chambers (Figure 6G). The gustatory test was performed as a two-bottle choice test between water and quinine solution. The CGRP^SPFp^ TetTox group consumed minimal quinine solution (0.5 mM), like controls, whereas the CGRP^PBel^ TetTox group showed much less aversion to quinine (Figure 6H).

These results indicate that both the CGRP^SPFp^ and CGRP^PBel^ neurons are necessary for the perception of innate sensory threat cues, as well as affective pain.

### Activating CGRP^SPFp^/CGRP^PBel^ to amygdala pathways induces negative valence

Next, we performed optogenetic gain-of-function experiments to test whether activation of these neurons is sufficient to induce negative affect in mice. We bilaterally injected AAV-DIO-ChR2 into the SPFp of *Calca^Cre^* mice and implanted optic fibers (200 μm, NA 0.22) above the injection site (Figure 7A). 20-Hz photo-stimulation of CGRP^SPFp^ neurons did not change responses in the hot plate thermal sensitivity test and the electronic von Frey mechanical threshold test for sensory and discriminative pain perception (Figures S8I-S8L). To test whether these neurons encode negative valence, we performed the real-time place aversion (RTPA) test. Optogenetic stimulation is delivered only when the test mouse stays on one side of a two-chamber apparatus (Stamatakis and Stuber, 2012; Figure 7B). Optogenetic activation of the CGRP^SPFp^ neurons induced aversion to the photo-stimulated chamber, suggesting that these neurons play a role in negative emotion or affective-motivational pain (Figures 7B and 7C). Next, we replaced the foot shock with photo-stimulation (20 Hz) as the unconditioned stimulus (US) in the context and cued fear conditioning test. This was to assess whether activation of CGRP^SPFp^ neurons was sufficient to induce fear behaviors. Context-dependent optogenetic conditioning was achieved by 10 mins of photo-stimulation in an open field arena; freezing behavior was then assessed in the same context 24 h after the conditioning (Figure S8M). The ChR2 group exhibited more freezing than the control group, suggesting that CGRP^SPFp^ activation can act as the US (Figure S8N). For cue-dependent optogenetic conditioning, photo-stimulation was associated with a tone as a non-noxious conditioned stimulus (CS+) in a fear conditioning chamber (Figure S8O). Both context and cue tests were performed after the conditioning. The ChR2 group exhibited more freezing in both context (Figure S8P) and cue tests (Figure S8Q). Together with the optogenetic conditioning results of CGRP^PBel^ neurons in the previous result (Han et al., 2015), these results indicate that activation of both CGRP^SPFp^ and CGRP^PBel^ neurons is sufficient to induce negative valence associated with affective-motivational pain perception.

**Figure 7.**
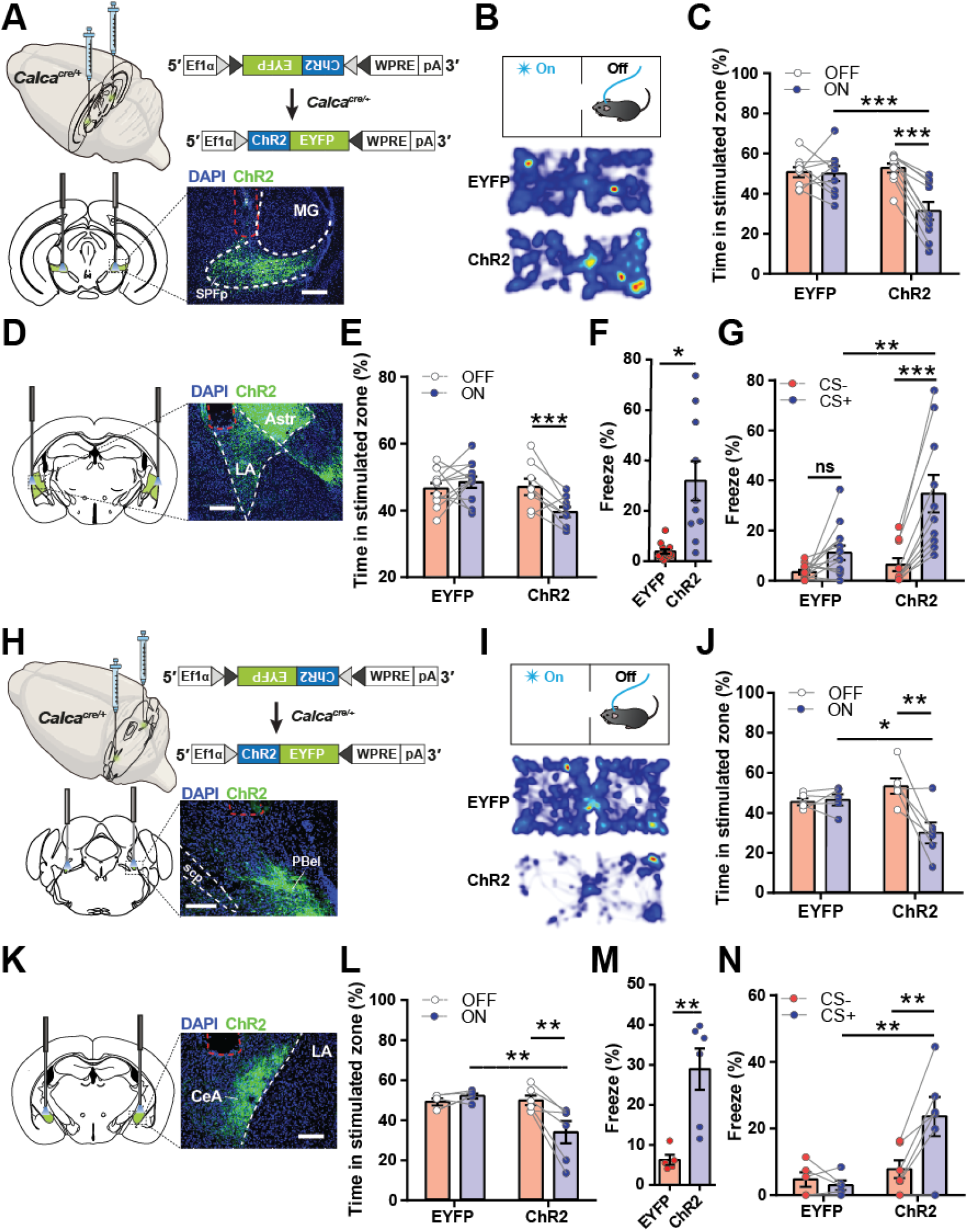
Activating the CGRP^SPFp→LA^ or CGRP^PBel→CeA^ pathways encodes negative valence. (**A**) Schematics and representative image of Cre-dependent expression of ChR2 in the SPFp of *Calca^Cre^* mice and optic fiber placement (red dotted line). (**B**) A schematic diagram and heatmap of the real-time place aversion (RTPA) test with CGRP^SPFp^ cell body stimulation. (**C**) Time spent in the stimulated zone during the RTPA test. (**D**) Representative image of optic fiber placement in the LA for terminal stimulation (red dotted line). (**E**) Time spent in the stimulated zone during the RTPA test with CGRP^SPFp→LA^ terminal stimulation.(**F**) Time spent freezing during the context test. (**G**) Time spent freezing during cue test after pairing optic stimulation and tone for cued fear conditioning. (**H**) Schematic and representative image of Cre-dependent expression of ChR2 in the PBel of *Calca^Cre^* mice and optic fiber placement (red dotted line). (**I**) Schematic diagram and heatmap of real-time place aversion (RTPA) test with CGRP^PBel^ cell body stimulation. (**J**) Time spent in the stimulated zone during the RTPA test with CGRP^PBel^ cell body stimulation.(**K**) Representative image of optic fiber placement in the CeA (red dot). (**L**) Time spent in the stimulated zone during the RTPA test with CGRP^PBel→CeA^ terminal stimulation. (**M**) Time spent freezing during the context test. (**N**) Time spent freezing during cue test after pairing optogenetic stimulation and tone for cued fear conditioning. Scale bars indicate 200 μm. **Statistics** (C) Repeated measure two-way ANOVA showed significance in laser x group interaction (F (1, 17) = 17.69, p = 0.0006) and laser (F (1, 17) = 20.19, p = 0.0003) but not group (F (1, 17) = 4.03, p = 0.0608). Laser ON and OFF in ChR2 (p < 0.0001), and EYFP and ChR2 during ON (p < 0.001) had statistically significant difference with Sidak’s multiple comparisons test. (E) Repeated measure two-way ANOVA showed significance in only laser x group interaction (F (1, 18) = 8.718, p = 0.0085) but not laser (F (1, 18) = 3.159 p = 0.0924) and group (F (1, 18) = 4.343, p = 0.0517). Laser ON vs OFF in ChR2 (p < 0.05), and EYFP vs ChR2 during ON (p < 0.01) have statistically significant difference with Sidak’s multiple comparisons test. (F) EYFP: 3.842 ± 0.89 (n = 13), ChR2: 31.92 ± 7.806 (n = 10). Unpaired t test (two-tailed), p = 0.0005. (G) Repeated measure two-way ANOVA showed significance in CS x group interaction (F (1, 21) = 12.41, p = 0.0020), CS (F (1, 21) = 38.26, p < 0.0001) and group (F (1, 21) = 8.392, p = 0.0086). CS-vs CS+ in ChR2 (p < 0.0001), and EYFP vs ChR2 during CS+ (p < 0.001) have statistically significant difference with Sidak’s multiple comparisons test. (J) Repeated measure two-way ANOVA showed significance in laser x group interaction (F (1, 9) = 10.49, p = 0.0102) and laser (F (1, 9) = 8.959, p = 0.0151) but not group (F (1, 9) = 1.159, p = 0.3096). Laser ON and OFF in ChR2 (p < 0.01), and EYFP and ChR2 during ON (p < 0.05) had statistically significant difference with Sidak’s multiple comparisons test. (L) Repeated measure two-way ANOVA showed significance in laser x group interaction (F (1, 9) = 12.91, p = 0.0058) and laser (F (1, 9) = 6.094, p = 0.0357) but not in group (F (1, 9) = 4.405, p = 0.0652). Laser ON and OFF in ChR2 (p < 0.01), and EYFP and ChR2 during ON (p < 0.01) had statistically significant difference with Sidak’s multiple comparisons test. (M) EYFP: 6.278 ± 1.244 (n = 5), ChR2: 28.93 ± 5.133 (n = 6). Unpaired t test (two-tailed), p = 0.0035. (N) Repeated measure two-way ANOVA showed significance in CS x group interaction (F (1, 9) = 12.67, p = 0.0061), CS (F (1, 9) = 8.226, p = 0.0185) and group (F (1, 9) = 6.314, p = 0.0332). CS-vs CS+ in ChR2 (p < 0.01), and EYFP vs ChR2 during CS+ (p < 0.01) have statistically significant difference with Sidak’s multiple comparisons test.

We then sought to characterize the functional downstream of the CGRP^SPFp^ neurons. To examine the functional connectivity of anatomical downstream regions from the CGRP^SPFp^ neurons, we performed *ex vivo* electrophysiology recording. AAV-DIO-ChR2-EYFP was injected into the SPFp of *Calca^Cre^* mice (Figure S9A). After four weeks of ChR2 expression, we performed whole-cell recordings of neurons from AStr, LA, and pIC to measure optogenetically-evoked excitatory/inhibitory postsynaptic currents (EPSC/IPSC; Figure S9B). We found that CGRP^SPFp^ neurons form functional glutamatergic synapses with neurons within AStr, LA (Figure S9C), and pIC (data not shown). Moreover, the onset of IPSCs lagged 4– 5 ms compared to the onset of EPSCs, indicating a feed-forward inhibition circuit. The number of cells that had both EPSCs and IPSCs, EPSCs only, IPSCs only, and non-responsive were also counted (Figures S9D-S9L). To investigate whether these connections form functional circuits that encode negative valence, we optogenetically stimulated axonal terminals from the CGRP^SPFp^ neurons and performed behavioral tests. AAV-DIO-ChR2-EYFP was injected into the SPFp of the Calca^*Cre*^ mice, and optical fibers were implanted into the postsynaptic areas of CGRP^SPFp^ neurons, namely the LA, AStr, and pIC (Figure 7D). Optogenetic activation of each of these three projections induced aversion in the RTPA experiment (Figures 7E and S9M), as observed in direct photo-stimulation of CGRP^SPFp^ cell bodies (Figure 7C). Cue-dependent optogenetic conditioning of the downstream circuits was then performed and only the CGRP^SPFp→LA^ circuit caused significant freezing in the context test (Figure 7F). The other two projections only showed a trend (Figure S9N). Increased freezing was observed in the cue test for all three projections (Figures 7G and S9O), but the most prominent effect was observed with the CGRP^SPFp→LA^ circuit.

The same behavioral experiments were performed as above with the CGRP^PBel^ neurons to compare their role in encoding negative valence with the CGRP^SPFp^ neurons. First, photo-stimulation of the CGRP^PBel^ neuronal cell body (Figure 7H) induced aversion during the RTPA test (Figures 7I and 7J). Then, we investigated the CGRP^PBel→CeA^ circuit as in Figure 7D-7H by optogenetic terminal stimulation (Figure 7K). Photo-stimulation of CGRP^PBel→CeA^ terminals induced aversion in the RTPA test (Figure 7L), and freezing in the optogenetic conditioning context, and cue tests (Figures 7M and 7N).

These results satisfy the idea that CGRP^SPFp→LA^ and CGRP^PBel→CeA^ circuits induce negative valence associated with either affective pain or innate sensory threat cues.

## Discussion

We report a genetically defined population of neurons that express the neuropeptide CGRP in the SPFp and PBel mediate perception of affective pain and innate sensory threat cues by relaying aversive sensory signals from the spinal cord and other sensory relay areas to the amygdala. These analyses provide the first evidence of convergent innate threat pathways that relay multisensory aversive sensory cues to the amygdala.

### CGRP neurons convey affective pain signals to the amygdala

Perception of pain protects us from physical harm by locating the source of a harmful stimulus. Painful experiences also elicit emotional and motivational responses, which help us remember these events and avoid similar stimuli in the future (Yeh et al., 2018). Thus, pain is not just a simple sensory process but also a complex cognitive process that generates sensory and emotional responses. This unique aspect of pain gives rise to the concept of two aspects of pain: sensory-discriminative and affective-motivational (Auvray et al., 2010; Melzack and Casey, 1968). It is thought that the sensory-discriminative aspect of pain is processed within the sensory cortex via the spino-thalamic tract (STT), and the affective-motivational aspect of pain is processed within the amygdala via the spino-parabrachial tract (SPT). Indeed, previous studies have shown that the PBN-to-CeA circuit is critical for affective-motivational pain perception (Han et al., 2015; Sato et al., 2015). However, it has also been suggested that the thalamus is actively involved in affective-motivational pain perception (Craig, 2003; Willis et al., 2002). Importantly, posterior regions of the thalamus (e.g., the VMpo in primates and humans, as well as the SPFp and PoT in rodents) are anatomically connected to limbic areas (Craig, 1998; Gauriau and Bernard, 2004a) and activated by noxious stimuli (Craig et al., 1994; Peschanski et al., 1981). Therefore, the thalamus likely also plays a critical role in affective-motivational pain perception. Our results demonstrate that a genetically defined population of neurons expressing the neuropeptide CGRP within the SPFp receives monosynaptic inputs from projection neurons within the spinal dorsal horn. They then project to specific nuclei within the amygdala, namely the AStr and LA (Figure 1). Multimodal nociceptive stimuli activate these neurons in an intensity-dependent manner in anesthetized mice (Figure 2). Inactivating these neurons attenuates the perception of affective-motivational pain, and pain signals induces the changes in synaptic plasticity of these neurons (Figure 3). These data indicate that CGRP^SPFp^ neurons create the spino-thalamo-amygdaloid affective pain pathway.

Unlike the STT, the SPT has been well-characterized as an affective-motivational pain pathway. Recent studies have shown that the lateral PBN receives direct nociceptive inputs from projection neurons within the spinal dorsal horn (Barik et al., 2020; Chiang et al., 2020; Choi et al., 2020; Deng et al., 2020). The dorsolateral PBN (PBdl) predominantly receives nociceptive inputs from the spinal cord and then projects to multiple limbic structures, such as the PAG, VMH, and ILN, thereby producing emotional and physiological changes in response to pain signals (Chiang et al., 2020; Deng et al., 2020). Although the PBdl does not directly project to the amygdala, it indirectly sends pain signals to the CeA through the PBel (Deng et al., 2020). In particular, CGRP^PBel^ neurons are critical for relaying aversive unconditioned stimuli to the CeA during aversive fear learning (Han et al., 2015). It has been shown that the dynorphin neurons in the PBdl project to the PBel (Chiang et al., 2020), and a recent study has shown that spinal projection neurons that express GPR83 directly innervate CGRP^PBel^ neurons to relay noxious signals (Choi et al., 2020). Therefore, it is clear that CGRP^PBel^ neurons relay nociceptive information from the PBdl and the spinal cord to the CeA. Our retrograde tracing results show that CGRP^PBel^ neurons receive less direct inputs from the spinal cord than the CGRP^SPFp^ neurons (Figure 1), and these neurons are activated slower than the CGRP^SPFp^ neurons (Figure S4C and S4J), implying that the CGRP^PBel^ neurons may receive nociceptive inputs more predominantly through the PBdl than through directly from the spinal cord. The CGRP^PBel^ neurons also exhibit intensity-dependent activation by multimodal nociceptive stimuli in anesthetized mice (Figure 2) and induce changes in synaptic plasticity by pain signals (Figure 3). Therefore, together with previous studies, our results reaffirm that CGRP^PBel^ neurons comprise the spino-parabrachio-amygdaloid pain pathway.

Comparing the CGRP^SPFp^ and CGRP^PBel^ neuronal activity in response to multimodal nociceptive stimuli at various intensities in anesthetized mice provides us with novel insights into understanding central affective pain pathways. First, unlike sensory pain signals that are conveyed to the contralateral side of the somatosensory cortex, these neurons are activated by both contralateral and ipsilateral noxious stimuli indicating that the affective pain pathway may not be strictly lateralized (Figure S4). Second, the CGRP^SPFp^ neurons more robustly respond to the mechanical stimulus, whereas the CGRP^PBel^ neurons respond more robustly to the thermal stimulus suggesting that these two parallel pathways may convey different modalities of nociceptive information (Figure 2). Lastly, the CGRP^PBel^ neurons are activated at lower stimulus intensity but respond slowly to nociceptive signals compared with the CGRP^SPFp^ neurons (Figures 2 and S4). Therefore, our results demonstrate that CGRP-expressing neurons in two brain areas (the SPFp and PBel) play complementary roles in relaying multimodal nociceptive signals from the spinal cord to the amygdala through two parallel ascending pain pathways, which are critical for affective-motivational pain perception.

### CGRP neurons convey multisensory innate threat cues to the amygdala

Both pain and innate sensory threats motivate animals to execute immediate avoidance behaviors to escape the threatening or tissue-damaging situation and produce long-lasting aversive memories. Indeed, Pavlovian threat conditioning uses noxious electric foot shock, an acute painful stimulus that motivates the animals to create aversive memory (LeDoux, 2012; Maren, 2001). However, the neural circuit mechanisms by which noxious information is conveyed to the amygdala during aversive learning is not fully understood. Recent advances in the neural circuit-based understanding of innate predator threat perception suggest that innate threat cues from each sensory modality are conveyed to discrete brain areas, such as the amygdala and hypothalamus through parallel pathways (Canteras, 2002; Gross and Canteras, 2012; Kunwar et al., 2015; Silva et al., 2013), which do not overlap with the unconditioned stimulus (affective pain) pathway in Pavlovian threat learning (Silva et al., 2016). However, it is beneficial for animals to use unified neural circuits that encode multimodal aversive sensory signals including pain because animals search for and detect imminent threats using multiple sensory modalities simultaneously. Moreover, previous clinical and animal studies have shown that pain and threat perceptions interact with each other (Berthier et al., 1988; Crook et al., 2014; Elman and Borsook, 2018; Lister et al., 2020). Therefore, it is plausible that there is a unified mechanism that conveys multisensory threat cues from multiple sensory modalities to the amygdala. Our results demonstrate that CGRP-expressing neurons in the PBel and the SPFp not only relay nociceptive stimuli to the amygdala during aversive learning, but also convey innate sensory threat cues from various sensory modalities (Figures 5 and 6). The CGRP^SPFp^ neurons relay aversive sensory cues from the somatosensory, visual, and auditory modalities to the LA, AStr, and pIC. By contrast, CGRP^PBel^ neurons relay aversive cues from all sensory modalities (somatosensory, visual, auditory, olfactory, and gustatory) (Figures 5, 6, and S3). In addition, previous studies have shown that CGRP^PBel^ neurons are activated by hypercapnic conditions (high CO_2_ levels) (Kaur et al., 2017; Yokota et al., 2015), and aversive visceral cues, such as lithium chloride and lipopolysaccharide (Carter et al., 2013; Paues et al., 2001). Therefore, it is tempting to speculate that CGRP^SPFp^ neurons relay exteroceptive threat signals, whereas CGRP^PBel^ neurons relay both exteroceptive and interoceptive threat cues to the amygdala.

Consistent with a previous report (Han et al., 2015), our data provide strong evidence that the CGRP^PBel^ neurons mediate aversive learning via their projection to the CeA (Figure 7K-7N). Our results seem to be contradictory to a recent study by Bowen et al., (2020), which argued that optogenetic stimulation of CGRP^PBel^ axon terminals in the VPMpc, instead of the CeA, evokes strong aversive memory. This discrepancy can be explained by the differences in optogenetic stimulation protocols. Whereas we used a 40 Hz light stimulation for 10 s as described (Han et al., 2015), Bowen et al. used a 30 Hz light train for only 2 s. Moreover, it is worth noting that axonal bundles from all glutamatergic neurons in the PBel that project to forebrain regions, including the CeA, BNST, and PSTN pass through the VPMpc (Huang et al., 2020), which suggests that optogenetic stimulation within the VPMpc activates both CGRP^PBel^ axonal terminals and axon bundles passing through this area. Therefore, an alternative explanation of their observation is that concurrent stimulation of all downstream areas produces stronger aversive memory than stimulating CeA alone.

Together, our results demonstrate that the US pathways for relaying electric foot shock cues during the Pavlovian fear learning also collectively relay innate threat cues from multiple sensory modalities.

### Pain-threat interactions in affective pain disorders

Our results show that the affective-motivational pain and multisensory threat stimuli arrive in the amygdala via the same neural pathways. Interestingly, pain-threat interactions have been reported in many human clinical cases. People with pain asymbolia, caused by damage to limbic brain areas, have the normal sensory perception of noxious stimuli, but they have impaired affective pain perception (Berthier et al., 1988; Rubins and Friedman, 1948). Interestingly, pain asymbolia patients often display deficits in perceiving general threats (Klein, 2015; Price, 2000), indicating that the perception of affective pain and other sensory threat cues share the same neural substrate. People with congenital insensitivity to pain (CIP) are insensitive to all sensory and affective components of pain, but they also display profound deficits in general threat perception, which is the primary cause of their short life expectancy (McMurray, 1955; Nagasako et al., 2003). CIP is caused by loss-of-function mutations in genes critical for pain transmission, such as *Scn9a* (Dabby, 2012; Fischer and Waxman, 2010; Lampert et al., 2010) and *Faah* (Drissi et al., 2020). *Scn9a* encodes voltage-gated sodium channel type 7 (Na_v_1.7), and *Faah* encodes fatty acid amide hydrolase, both of which are critical for pain transmission in the spinal cord (Cajanus et al., 2016; Kim et al., 2006; Nantermet and Henze, 2011; Nassar et al., 2004). However, the functional loss of these genes in the spinal neurons cannot explain the insensitivity to general threats exhibited by CIP patients. Surprisingly, our cell type-specific transcriptome analysis revealed that *Scn9a* and *Faah* transcripts are highly enriched in both CGRP^PBel^ and CGRP^SPFp^ neurons (Figure 4). Thus, mutations in these genes may prevent these neurons from relaying sensory threat signals to the amygdala, thereby causing insensitivity to general threats in CIP. This speculation should be addressed by testing the causal relationship between mutations in these genes in CGRP neurons and threat perception.

Opposite clinical cases also exist. People with affective pain disorders, such as migraine, and fibromyalgia experience chronic pain and suffer from hypersensitivity to normal sensory stimuli (Demarquay and Mauguière, 2016; Harriott and Schwedt, 2014; Harte et al., 2016; López-Solà et al., 2017). Further, normal sensory stimuli often trigger or aggravate their pain symptoms (Bar-Shalita and Cermak, 2020; Bar-Shalita et al., 2019). Surprisingly, the gene *Cacna1a*, which has been linked to migraines, is highly enriched in both CGRP^PBel^ and CGRP^SPFp^ neurons (Figure 4). Further studies should address the causal relationship between loss of *Cacna1a* function in CGRP neurons and sensory hypersensitivity in migraine. In addition, CGRP signaling is a proven therapeutic target for treating migraine (Ashina, 2020). Therefore, we speculate that the CGRP-expressing neurons characterized in this study may serve as the functional substrate for sensory hypersensitivity in migraine. Identifying membrane proteins commonly expressed in both the CGRP^PBel^ and CGRP^SPFp^ neurons (Figure 4C) may provide potential candidates of therapeutic targets for treating affective pain-, and innate threat-related disorders, such as migraine, fibromyalgia, phobias, panic disorder, and post-traumatic stress disorder. Indeed, the CGRP receptor antagonists or neutralizing monoclonal antibodies are promising drugs for treating migraine, but they may also serve as potential therapeutic interventions for treating the disorders mentioned above.

## Conclusion

Our findings demonstrate that the CGRP^SPFp^ neurons create a novel spino-thalamo-amygdaloid affective pain pathway and, together with the previously characterized CGRP^PBel^ neurons, serve as complementary parallel pathways for conveying the unconditioned stimulus during Pavlovian threat learning, as well as multisensory innate threat cues from all sensory modalities to the amygdala (Figure S10). The discovery of a unified threat perception system that conveys multimodal interoceptive and exteroceptive aversive sensory stimuli greatly enhances our understanding of the neural mechanisms of innate threat perception. These insights also provide opportunities to discover novel targets for developing therapeutic interventions against affective pain- and innate threat-related disorders.

## Acknowledgments

We thank Dr. D. O’Keefe, Ms. C. Jia, and Han lab members for critical discussions during manuscript preparation. S.H. is supported by 1R01MH116203 from NIMH and the Bridge to Independence award from the Simons Foundation Autism Research Initiative (SFARI #388708). S.L. is supported by the Salk Women & Science Special Award, the Mary K. Chapman Foundation, and the Jesse & Caryl Philips Foundation.

## Author Contributions

S.H. conceived of the idea and secured funding. S.H., S.J.K., S.L., and M.Y. designed and performed the experiments. S.H. and S.J.K wrote the manuscript. S.J.K. performed most of the CGRP^SPFp^ experiments. S.L., and S.J.K. performed CGRP^PBel^ experiments. M.Y. performed electrophysiology and CGRP^PBel^ RTPA. D.I.K., and J.H.K. performed RiboTag experiments, and T.G.O analyzed it. R.M.E. provided resources for Ribotag analysis. J.P. performed spinal projection histology experiments. M.G. provided *Cdx2^FlpO^* mouse line. K.F.L. provided resources for microscopy.

## Supplementary Information

### Materials and methods

#### Animals

All protocols for animal experiments were approved by the IACUC of the Salk Institute for Biological Studies according to NIH guidelines for animal experimentation. The *Calca^Cre^*, and *Tacr1^Cre^* transgenic mouse lines used in this study were generated from the Richard Palmiter’s laboratory (Han et al., 2015 or Carter et al., 2013) *Cdx2^FlpO^* line was generated from the Martyn Goulding’s laboratory. *Calca^CreER^* mouse line was generated from the Pao-Tien Chuang’s laboratory. RiboTag *Rpl22^HA/HA^* (Stock No. 011029) and Ai65 (Stock No: 021875) mouse line was obtained from the Jackson Laboratory. All mouse lines are backcrossed with C57Bl/6J for > 6 generations. Male and female mice were used in all studies. Animals were randomized to experimental groups and no sex differences were noted. Mice were maintained on a normal 12 hours light/dark cycle and provided with food and water *ad libitum*.

#### Stereotaxic surgery for virus injection and optic fiber implantation

Mice were anesthetized by isoflurane gas anesthesia (induction at 3.5%, and maintenance at 1.5-2%, the Dräger Vapor^®^ 2000; Draeger, Inc., USA). Mice were then placed on a stereotaxic frame (David Kopf Instruments, USA). Holes were drilled with a micromotor handpiece drill (Foredom, USA) after the exposure of the skull. The virus was injected using a syringe (65458-01, Hamilton, USA) connected to an ultra-micropump (UMP-3, World Precision Instruments, USA). Unilateral (right side) and bilateral injections were made for the following target regions: SPFp (antero-posterior (AP), −3.1 mm; medio-lateral (ML), 2.0 mm; dorso-ventral (DV) −3.6 mm from bregma) or PBel (AP, −5.1 mm; ML, 1.35 mm; DV, −3.5 mm). Viruses were injected at a rate of 0.08 μl/min (total volume of 0.75 μl for optogenetic projection studies and 0.5 μl for all the others) and the syringe needle was slowly removed from the injection site seven-minute after injection. To determine the inputs to CGRP^SPFp^ and CGRP^PBel^ neurons, 0.5 μl of AAV8-hSyn-FLEX-TVA-P2A-GFP-2A-oG (Salk Institute viral vector core, USA) was injected into the SPFp or PBel of *Calca^Cre^* transgenic mice. After three weeks, 0.5 μl of EnvA-ΔG-rabies-mCherry (Salk Institute viral vector core, USA) was injected, and the mice were sacrificed five days after the injection. To silence the CGRP^SPFp^ and CGRP^PBel^ neurons, 0.5 μl of AAV-DIO-TetTox-GFP or AAV-DIO-EYFP was injected into the SPFp or PBel of *Calca^Cre^* transgenic mice, and experiments were performed two weeks after injection. For fiber photometry experiments, mice were injected with 0.5 μl of either AAV-DIO-GCaMP6m, AAV-DIO-GCaMP7s or AAV-DIO-EYFP into the SPFp or PBel of *Calca^Cre^* mice. Stainless-steel mono fiberoptic cannulas (400 um diameter, 0.37 NA, Doric Lenses) were implanted above the SPFp or PBel. For electrophysiology, mice were injected with 0.5 or 0.75 μl of AAV1-DIO-ChR2-EYFP into the SPFp or PBel of *Calca^Cre^* mice. Experiments were performed two weeks after viral injection for recording SPFp and PBel neurons or four weeks after injection for recording cells in terminal regions. For optogenetics, mice were injected with 0.5 or 0.75 μl of AAV1-DIO-ChR2-EYFP or AAV-DIO-EYFP into the SPFp or PBel of *Calca^Cre^* mice and custom made mono fiberoptic cannula (200 um diameter, 0.22 NA) were implanted above SPFp (0.5 mm above the injection site), PBel (0.5 mm above the injection site), Astr (AP, −1.8 mm; ML, 3.3 mm, DV, −3.8 mm from bregma), lateral amygdala (LA; AP, −1.8 mm; ML, 3.6 mm, DV, −4.0 mm from bregma), pIC (AP, −1.5 mm; ML, 4.6 mm, DV, −3.0 mm from bregma) or central amygdala (CeA; AP, 1.2 mm; ML, 2.7 mm; DV, −4.2 mm from bregma). Experiments were executed two weeks after injection to manipulate SPFp neurons or four weeks later for terminal stimulations.

#### Histology and quantification of rabies tracing experiment

Mice were intracardially perfused with 4% paraformaldehyde in PBS 5 days after the rabies virus injection. Spinal cords were post-fixed at 4°C for 1 h and dehydrated with 30% sucrose at 4°C overnight. 40 um transverse sections were obtained with a cryostat (CM 1950, Leica, USA) throughout the spinal cord. Spinal cord slices were directly dry mounted on superfrost plus microscope slide glasses (12-550-15, Fisher Scientific, USA). The labeled neurons were counted manually by dividing the transverse spinal sections into four groups (cervical, thoracic, lumbar, and sacral) or different dorsal horn layers. Brains were kept in 4% PFA overnight for post-fixation and dehydrated in 30% sucrose for 1-2 days before sectioning. Frozen brains were cut into 50 μm coronal slices with a cryostat and stored in Phosphate buffered saline before mounting. Both spinal cord and brain tissues were mounted on a slide glass with a DAPI containing mounting solution (0100-20, SouthernBiotech, USA).

#### Fiber photometry

Bulk calcium signals from the CGRP^SPFp^ and CGRP^PBel^ neurons were monitored using a custom-built fiber photometry system based on the open-source pyPhotometry platform (https://pyphotometry.readthedocs.io/en/latest/). 465 nm LED was used to induce Ca^2+^ dependent fluorescence signals, and 405 nm LED was used for Ca^2+^ independent (isosbestic) fluorescence signals. Motion corrected ΔF/F was calculated by a post-hoc analysis (ΔF/F = F465-F405fit / F405fit). The least-squares polynomial function was used to calculate F405fit, and the area under the curve was used to analyze the data.

#### Multi-modal aversive stimuli experiments

##### Mechanical and thermal stimuli

Mechanical and thermal stimuli were applied to mice forepaw, hind paw, or tail. Mechanical pressure was applied using a dial tension gauge (ATG-300-1, Vetus Instruments, USA) with stimulus strength 0, 50, 100, 200 g. The thermal stimulus was applied using a custom-made temperature-controlled hot-rod (TA4-SNR+K, Mypin, China) at 25, 35, 45, and 55°C. A stable baseline was recorded first for 10 s, and stimuli were applied immediately after 5 seconds.

##### Formalin test

For fiber photometry, lightly anesthetized mice were placed in the stereotaxic frame head fixed to minimize movement. 10 μl of 4% formalin (1.6% Paraformaldehyde, 19210, Electron Microscopy Sciences, USA) was injected subcutaneously on the contralateral forepaw after at least 5 min of stable baseline. Calcium transients were recorded for 45 min. For the loss of function experiment, 10 μl of 4 % formalin was injected subcutaneously on one side of the forepaw. Mice were then placed in a Plexiglas chamber (10 x 10 x 13 cm) with a mirror placed behind. Behaviors were recorded for an hour, and licking behaviors were manually counted throughout the experiment.

##### Auditory and visual stimuli

For both auditory and visual stimuli experiments, mice were placed in a cylinder-shaped arena (11 cm diameter with 15 cm height) with homecage bedding and were habituated for 30-120 min. For auditory experiments, after a stable 10s baseline, an intense sound (85 dB, 2 s) or a control sound (70 dB) was played. For the loss-of-function experiments, mice were placed inside an open field chamber and were habituated for 10 min. After 1 min baseline, an intense sound (85 dB, 2 s) was delivered three times with an inter-stimulus interval of 28 s. All the trials were recorded by a USB camera (DFK 33GX236, Imagine Source, Germany) attached to a computer, and freezing behavior was analyzed using video-tracking software (Ethovision XT, Noldus, Netherlands). For visual looming experiments, after a stable 10 s baseline, an expanding looming stimulus (2 s) was delivered three times with 10 s inter-stimulus interval with a LED screen facing the arena from above. For the loss-of-function experiment, mice were placed in a cage with bedding and were positioned under the same LED screen. Mice were habituated for 20-30 min. When mice were in the center, the expanding looming stimulus (2 s) was delivered three times with 10 s inter-stimulus interval. All the trials were recorded by a USB camera (DFK 33GX236, Imagine Source, Germany) attached to a computer, and freezing behavior was analyzed using video-tracking software (Ethovision XT, Noldus, Netherlands) with manual counting for the duration of tail rattling behaviors.

##### Gustatory stimulus

For fiber photometry, mice were placed in the same arena for auditory and visual stimulus experiments with an additional 2 cm drilled hole. The water bottle spout was inserted into the hole, and the calcium signal was measured when the mice were licking. The bottle was filled with water or quinine (0.5 mM, QU109, Spectrum Chemical, USA). For the loss-of-function experiment, mice were water-deprived overnight. The next day, mice were placed in a homecage with a water-, and 0.5 mM quinine-containing bottle inserted into the water valve slot. Mice were allowed to drink for 10 min without habituation. All the trials were recorded by a USB camera (DFK 33GX236, Imagine Source, Germany) attached to a computer, and the licking behaviors were counted manually.

##### Olfactory stimulus

For fiber photometry, mice were placed in the same arena for gustatory stimulus experiments. Water-or Trimethylthiazoline (TMT, 97%, 5 μl, 1G-TMT-97, BioSRQ, USA)-soaked cotton swap was introduced into the hole. Calcium signals were measured when mice smelled the cotton swap. For the loss-of-function experiment, mice movement was tracked in a two-chamber arena (30 x 60 x 30 cm) with a USB camera (DFK 33GX236, Imagine Source, Germany) using video-tracking software (EthoVision XT 12, Noldus, Netherlands). Two Petri dishes with small holes were placed in each chamber (one at the corner of the left chamber, and the other to the corner of the right chamber). On day 1, mice were able to habituate and explore the arena for 10 min. The next day, a water-soaked cotton swap, or TMT-soaked cotton swap were placed in each dish. Mice were first placed at the center and monitored for 10 minutes as they interreacted with the two dishes.

##### Foot shock

A fear-conditioning chamber (26 x 30 x 33 cm, ENV-007CT, MED Associates INC, USA) with a metal grid floor (ENV-005, MED Associates INC, USA) connected to a standalone aversive electric shock stimulator (ENV-414S, MED Associates INC, USA) was used for foot shock delivery. A USB camera (DFK 33GX236, Imagine Source, Germany) was connected to a computer, and the video tracking software (Ethovision XT, Noldus, Netherlands) was used for shock delivery and behavioral analysis. The chamber was enclosed in a light- and sound-attenuating cubicle (ENV-018MD, MED Associates INC, USA). The chamber was cleaned with 70% ethanol and double distilled water between each trial.

For fiber photometry and the loss-of-function experiment, mice were placed inside the chamber without habituation. After 2 min of baseline, an electric shock (2 s, 0.6 mA) was delivered, and the behavior was recorded for an extra 2 min. Freezing behavior was monitored before (habituation), after (conditioning), and one day after (post-test) the shock.

##### Elevated plus maze test

A custom-built elevated plus maze with two transparent closed arms (77 x 7 x 30 cm) and two open arms (77 x 7 x 2 cm) was used to monitor the anxiety-like behaviors of test mice. This maze was elevated 70 cm above ground for all tests. Mice were placed to the tip of the open arm by facing towards the center of the maze. The behavior was video recorded for 10 min and tracked with a video-tracking software (EthoVision XT 12, Noldus, Netherlands). Both 70% ethanol solution and deionized water were used to clean the maze immediately after each trial.

##### Hot plate test

Mice were placed into a cylinder-shaped transparent Plexiglas chamber (11 cm diameter with 15 cm length) on a heated hot plate (48 or 55°C, PE34, IITC Life Science, USA). The latency of various pain responses (hind paw shake, lick, or jump) was measured manually.

##### Electronic von Frey test

A Dynamic Plantar Aesthesiometer (37450, Ugo Basile, Italy) was used to measure the mechanical pain thresholds. Mice were placed inside a Plexiglas chamber (10 x 10 x 13 cm) on a metal mesh floor and were habituated for 30 min. The max force of the system was set to reach 50 g at 20 s. The blunt metal rod of the aesthesiometer was placed under the hind paw and gradually protruded as the mice were immobile but awake. The latency and force delivered were automatically recorded as the mouse withdraw hind paw from the metal rod. The measurement was performed 5 times with 5-10 min intervals in between trials and averaged for a final mechanical threshold value.

#### Optogenetics

A 470 nm laser (LRD-0470-PFFD-00100-05, LaserGlow Tech., Canada) was used for all optogenetic experiments in this study. Optic fibers were bilaterally connected to pre-implanted optic ferrules on the mice. All mice were optogenetically stimulated 90 min before sacrifice for cFos immunohistochemistry.

##### Hot plate test

The experiments were performed as described in the ‘*Hot plate test’* section above with minor modification for optogenetic stimulation. The order of laser ON or OFF was counterbalanced, and the interval time between each experiment was more than 30 min.

##### Electronic von Frey test

The experiments were performed as described above in the ‘*Electronic von Frey test’* with minor modification for optogenetic stimulation. The laser was on immediately before the metal rod touched the paw pad and turned off right after paw withdrawal.

##### Real-time place aversion (RTPA)

A two-chamber arena (30 x 60 x 30) was used for the RTPA test. The behavior was tracked with a USB camera (DFK 33GX236, Imagine Source, Germany) using video-tracking software (EthoVision XT 12, Noldus, Netherlands). After connecting the optic fiber, mice were placed in one side of the chamber. No stimulation was given for 10 min baseline. Afterward, one side of the chamber was randomly selected, and the mouse was photostimulated (20 Hz for cell body stimulation, and 40 Hz for terminal stimulation, 8-9 mW) for 20 min. The stimulated side was counterbalanced between animals. Mice showing over 15% preference to one side during baseline were excluded.

##### Context-dependent optogenetic conditioning

An open field arena (40 x 40 x 30 cm) was used for context-dependent threat conditioning. After 10-min habituation to head-attached optic fibers in the home cage, mice were placed in the novel open field area and received photostimulation (20 Hz, and 8-9 mW) throughout the experiment. After 24 h, the mouse was re-introduced in the same context to test whether the photo-stimulation produced aversive memory. All the trials were recorded by a USB camera (DFK 33GX236, Imagine Source, Germany) attached to a computer, and freezing behavior was analyzed by a video-tracking software (EthoVision XT 12, Noldus, Netherlands).

##### Auditory cue dependent optogenetic conditioning

The same fear-conditioning chamber and the settings as described in the ‘*Foot shock’* section above were used. Two speakers (AX210, Dell, USA) were placed beside the chamber for CS. On day 1, the test mouse was habituated with the conditioning chamber, which was cleaned with 70% ethanol and DW immediately after each test. During habituation, optic fibers were connected bilaterally to the optic ferrules on the mouse’s head, and the CS+ (30 s, 3 kHz pure tone, 75 dB) was delivered to the test mouse six times with random inter-event intervals. On day 2, the test mouse was returned to the same context with optic fibers connected and received 10-s photostimulation (20 Hz frequency for cell body and 40Hz for terminal stimulation, 8-9 mW intensity) as the US, which was co-terminated with CS+ six times with random inter-event intervals. On day 3, the conditioned mouse without the optic fiber connected was returned to the same context for 2 min. On day 4, a conditioned mouse without the optic fiber connected was introduced to a new context (a glass cylinder wrapped with a non-transparent material), and the CS+ was delivered without the US. All the trials were recorded by a USB camera (DFK 33GX236, Imagine Source, Germany) attached to the computer, and freezing behavior was analyzed by a video-tracking software (EthoVision XT 12, Noldus, Netherlands).

#### Immunohistochemistry

Mice were perfused intracardially with % PFA solution in PBS, and the brain was extracted. The brain was kept in 4% PFA overnight for post-fixation and dehydrated in 30 % sucrose for 1-2 days before sectioning. Frozen brains were cut into 40 μm coronal slices with a cryostat (CM 1950, Leica, USA) and washed with PBST (Phosphate buffered saline with 0.1% Tween-20 (BP337-500, Fisher BioReagents, USA)). Initial blocking was performed by 1hr incubation with 3% normal donkey serum (NDS, 017-000-121, Jackson ImmunoResearch Laboratories, Inc., USA). After another round of washing with PBST, the slices were incubated with anti-GFP (diluted 1:100 in 3% NDS, GFP-1020, Aves, USA), anti-fos (1:10000, rabbit polyclonal), anti-Na_v_1.7 (1:200, ASC-008, Alomone), anti-Ca_v_2.1 (1:100, ACC-001, Alomone, Isreal), or anti-FAAH (1:250, 101600, Cayman, USA) antibody at 4 °C overnight. The next day, brain tissues were rinsed with PBST, then incubated with anti-rabbit Alexa Fluor^®^ 647-secondary antibody (1:500, 711-605-152, Jackson ImmunoResearch Laboratories, Inc., USA), and / or anti-chicken Alexa Fluor^®^ 488-secondary antibody (1:500, Jackson ImmunoResearch Laboratories) for 1 h. After washing these brain slices with PBS, they were mounted on slide glass (12-550-143, Fisher Scientific, USA) with DAPI containing mounting solution.

#### Preparation of acute brain slices and electrophysiology

Mice were anesthetized with isoflurane and perfused via the vascular system using ice-cold cutting solution (110.0 mM choline chloride, 25.0 mM NaHCO_3_, 1.25 mM NaH_2_PO_4_, 2.5 mM KCl, 0.5 mM CaCl_2_, 7.0 mM MgCl_2_, 25.0 mM glucose, 5.0 mM ascorbic acid and 3.0 mM pyruvic acid, bubbled with 95% O_2_ and % CO_2_). After decapitation, brains were quickly removed and chilled in an ice-cold cutting solution. Coronal slices containing the SPFp, PBel (250 μm) or the amygdaloid complex (300 μm) were cut by using a Leica VT 1200S Vibratome (Leica Biosystems Inc.), and subsequently transferred to a storage chamber containing artificial cerebrospinal fluid (aCSF; 124 mM NaCl, 2.5 mM KCl, 26.2 mM NaHCO_3_, 1.2 mM NaH_2_PO_4_, 13 mM glucose, 2 mM MgSO_4_ and 2 mM CaCl_2_, at 32 °C, pH 7.4, bubbled with 95% O_2_ and 5% CO_2_). After at least 30 min recovery time, slices were transferred to room temperature (22–24°C) for at least 60 min before use. Slices were transferred into the recording chamber, perfused with aCSF (flow rate around 2 ml/ min). The temperature of aACSF was held constant at 32°C by TC-324C temperature controller (Warner Instruments). Since CGRP-positive neurons express EGFP under the *Calca* promoter, they were visualized under Scientifica Microscope equipped with epifluorescence illumination at 490 nm LED. The Astr, LA, and IC neurons were visualized under trans-illumination. Whole-Cell patch clamp was performed with Multiclamp 700B amplifiers (Molecular Devices). Signals were digitized at 10 kHz with Digidata 1550B (Molecular Devices). For evoked EPSCs, synaptic responses were evoked with a broken glass pipette positioned 100 μm away from the recording glass electrode (3.0~5.0 MOm, back filled with internal solution: CsMeSO_3_ 130 mM, CsCl 5, HEPES 10 mM, MgCl_2_ 2.5 mM, EGTA 0.6 mM, Sodium phosphocreatine 10 mM, Na_2_ATP 4 mM and Na_3_GTP 0.4 mM, pH 7.23, 285 Osm). The stimulus was given at 0.1 Hz. AMPA EPSC was recorded holding at −70 mV for 10 to 30 sweeps to get a stable response. NMDA EPSC was recorded at +40 mV for 10 – 15 sweeps. To ensure that the EPSCs were stable, the holding potential was set to −70 mv to check the AMPA EPSC change after NMDA EPSC recording. Evoked EPSCs were recorded with picrotoxin (100 μM) in the aCSF. mEPSCs were recorded in the presence of tetrodotoxin (1 μM) and picrotoxin (100 μM). To record optogenetically evoked EPSC and IPSC, slices were harvested from the AAV-DIO-ChR2-EYFP injected *Calca^Cre^* mice brain. 2-ms 470 nm LED light (TTL from Clampex to Cool Led pE-300) was illuminated through 40X NA 0.8 objective lens at 0.1 Hz to evoke optogenetically evoked postsynaptic current. The internal solution was calculated to make chloride reversal potential at −70 mV. EPSCs were recorded at −70 mV, and IPSCs were recorded at 0mV. CNQX (10 μM) was perfused to check the glutamatergic synapse. EPSCs and IPSCs were analyzed using pCLAMP 10 software (Molecular Devices). NMDA EPSCs were defined as signals 100 ms apart from stimulus artifacts. mEPSCs were analyzed using Mini Analysis Program (Synaptosoft).

#### Imaging

The images were taken with an automatic fluorescence microscope (BZ-X710, Keyence, USA) using included imaging software (BZ-X viewer, Keyence, USA) or with a scanning confocal microscope (FV 1000, Olympus, Japan) using with Fluoview software (Olympus, Japan). For quantification purposes, images were processed with the same gain, offset, and exposure time. Cell counting for retrograde tracing was done manually.

#### RiboTag Transcriptomic Profiling

To label the active transcriptome of CGRP^SPFp^ and CGRP^PBel^ neurons, we crossed *Calca^CreER^* with RiboTag *Rpl22^HA/HA^* mice. To induce gene expression, 200 μl of tamoxifen freshly prepared with 20 mg/ml in corn oil and dissolved overnight with continuous agitation was administered intraperitoneally for five consecutive days in each mouse. Experiments were performed two weeks after the final tamoxifen injection. 250 μm thick slices containing the PBel and the SPFp were obtained using a VT 1200S Vibratome (Leica, Germany). The region of interest was further dissected using surgical scissors under the stereoscope. Tissues of interest from four *Calca^CreER^*; RiboTag crossed mice (10-12 weeks old) were collected into 1.5 mL microcentrifuge tubes containing homogenization buffer and were mechanically dissociated and lysed using pellet pestles (Cat.no.7495211500-DS, DWK Life Sciences LLC, USA). Total RNA was extracted from 15% of cleared lysate for input samples. The remaining lysate was incubated with mouse anti-HA antibody (Cat.no.MMS-101R, Biolegend, USA) and was rocked for 4 hours at 4 °C. Afterward, magnetic beads (Cat.no.88803, Thermo Fisher Scientific, USA) were added, and the solution was incubated overnight, rocking at 4 °C. The beads were washed three times in high salt solution. The bound ribosomes and RNA were separated from the beads by 30 s of vortexing in RLT lysis buffer as IP. All RNA samples were purified from the IP and corresponding input samples (Qiagen RNeasy Mini Kit, cat.no. 74104), then quantified with the Qubit RNA Assay Kit (Invitrogen, USA) and analyzed with the RNA 6000 Pico Kit (Agilent, USA). Isolated RNA was prepared using the Trio RNA-Seq (Cat. No. 0507-08; NuGEN, USA). Briefly, cDNA was synthesized from the total RNA using reverse transcriptase with oligo dT and resynthesized to produce double-stranded cDNA. After amplifying double-stranded cDNA, cDNA was purified with AMPure XP Beads (Cat. No. A63881; Beckman Coulter, USA), fragmented to the library, and classified using a barcoded adaptor. All libraries were quantified by qPCR and analyzed with the RNA 6000 Pico Kit. RNA library quality was checked using the 2100 Bioanalyzer (Agilent, USA). Barcoded samples were pooled and sequenced on the NextSeq500 (Illumina, USA) with the 75 bp read length single-end. Image analysis and base calling were conducted using the Illumina CASAVA-1.8.2 software. The FastQC package was utilized to evaluate the sequencing read quality. Fastq reads were then aligned to the reference genome (GRCm38.p6) using the STAR tool (version 2.7.2) in a pair-end mode. The quantification package RSEM (version 1.2.28) was employed to calculate gene expression from BAM files using the default setting changed to pair-end mode. In doing so, estimated count and TPM (Transcripts Per Million) were generated. Fold changes were calculated from TPM values (estimated counts, > 20) between HA-tag and HA negative controls., The ggplot2 package from R was utilized to visualize fold changes. UP (> 2.5-fold change) and DOWN (< −2.5-fold change) were highlighted with orange and blue colors, respectively.

#### Statistical analysis

All data are shown as mean ± s.e.m. and analyzed using Student’s t-test, one-way ANOVA with Tukey’s post hoc comparison, and two-way ANOVA with Sidak’s post hoc comparison. All the statistical analyses were done using Prism 6 (GraphPad Software Inc., USA). NS p>0.05, * p < 0.05, ** p < 0.01, *** p < 0.001

## Supplementary Figure Legends

**Figure S1.**
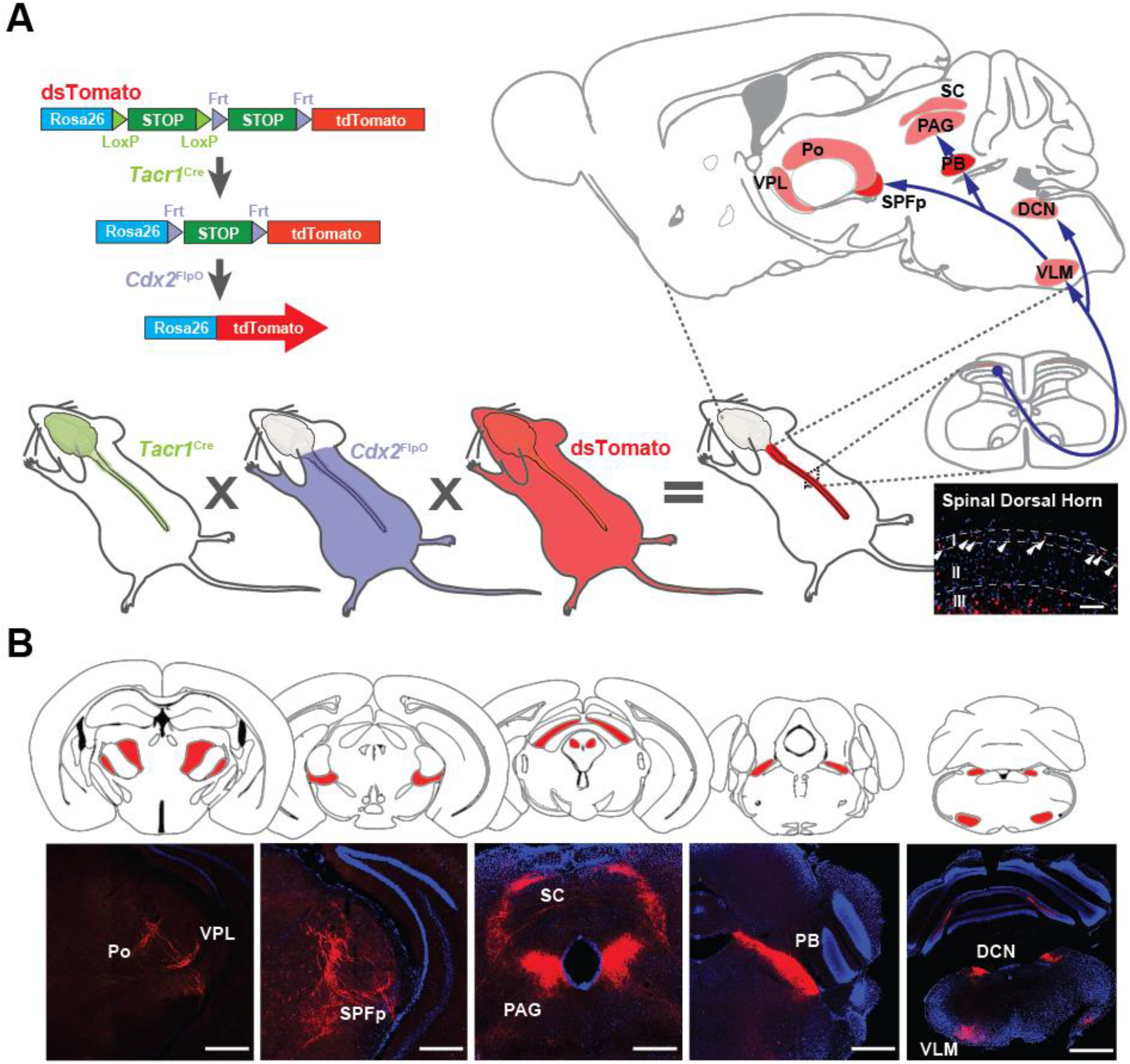
Identification of direct spino-recipient areas in the brain by specific genetic labeling of *Tacr1*-expressing spinal projection neurons. (**A**) Schematics of the triple cross strategy to specifically label *Tacr1*-expressing neurons in the spinal dorsal horn. Scale bar indicates 200 μm. (**B**) Spinal *Tacr1*-expressing neurons send projections to the posterior complex of the thalamus (PO), ventral posterolateral nucleus of the thalamus (VPL), the ventral posteromedial nucleus of the thalamus (VPM), SPFp, superior colliculus (SC), periductal gray (PAG), PB, dorsal column nuclei (DCN) and ventrolateral medulla (VLM). Scale bar indicates 500 μm.

**Figure S2.**
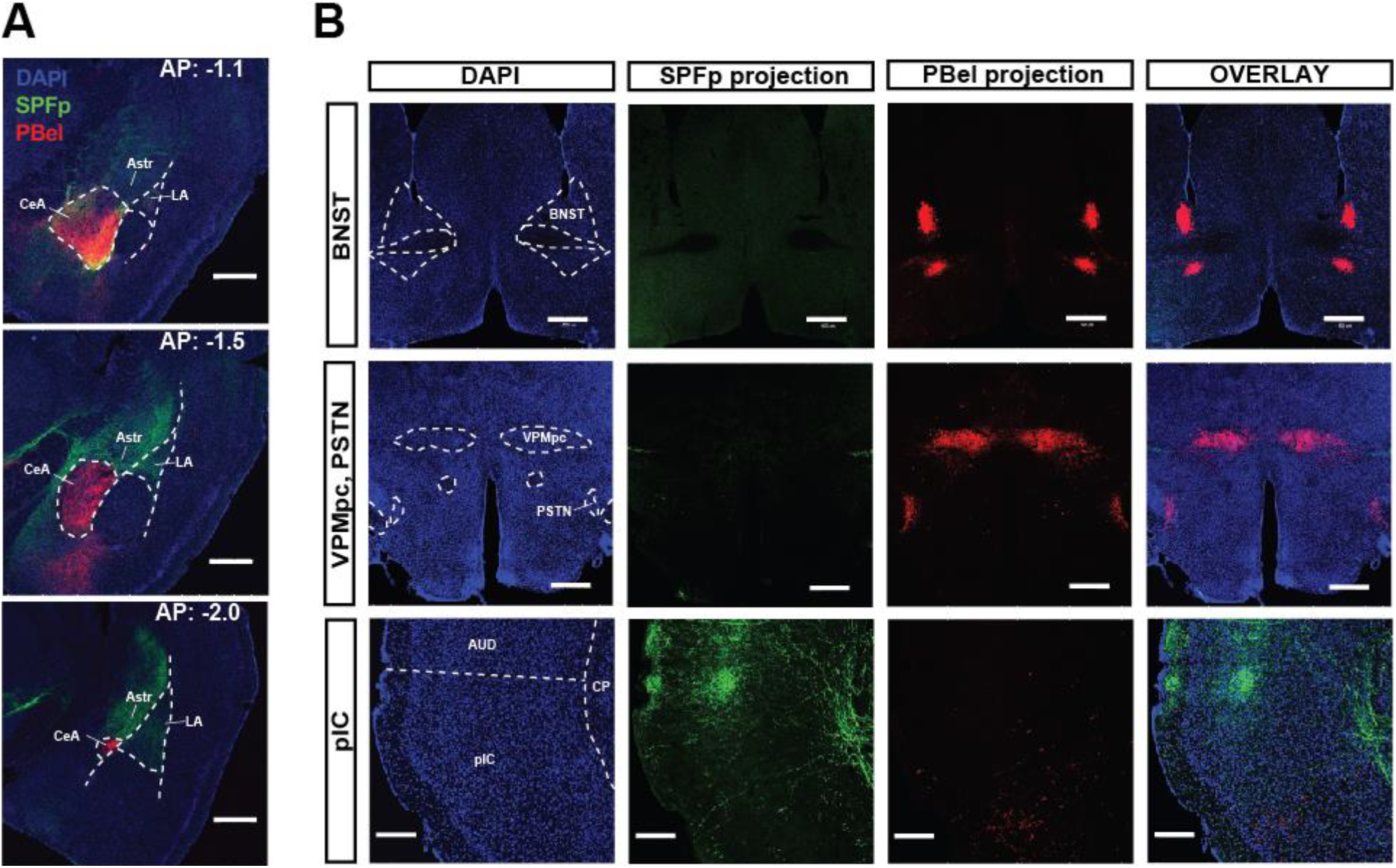
Projections from CGRP^SPFp^ and CGRP^PBel^ neurons. (**A**) Additional images from Figure 1A, B. Projections are prominent in the amygdala regions and have distinct patterns along the anterior-posterior axis; AP: −1.1, −1.5 and −2.0 mm from bregma. Scale bar indicates 500 μm. (**B**) The CGRP^PBel^ neurons also project to BNST, VPMpc, PSTN, and ventral pIC. The CGRP^SPFp^ neurons project to the auditory cortex, and dorsal pIC. Scale bars indicate 500 μm for BNST, and VPMpc; 200 μm for pIC.

**Figure S3.**
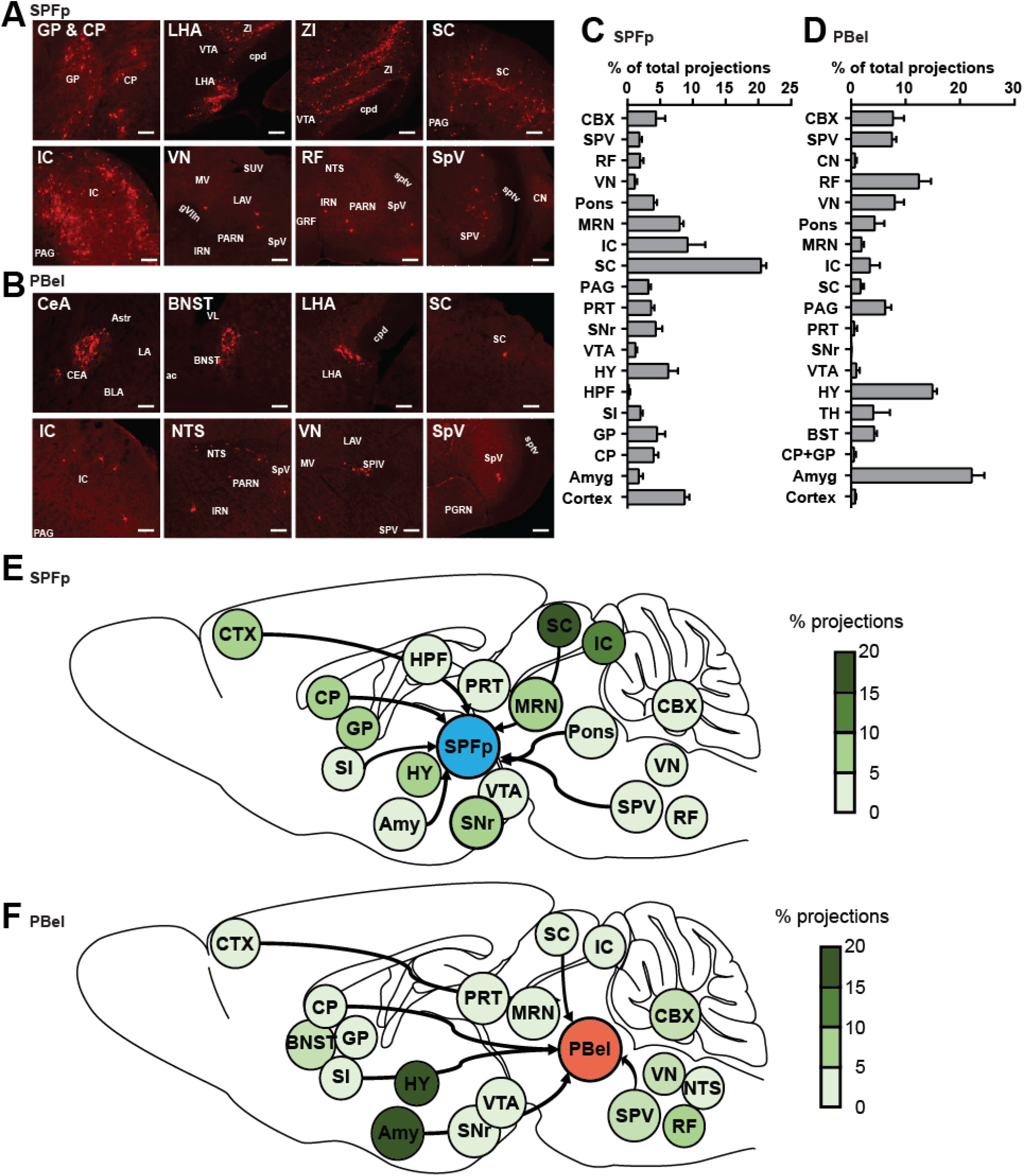
Retrograde tracing from CGRP^SPFp^ and CGRP^PBel^ neurons. (**A** and **B**) Example schematic of brain regions that send inputs to CGRP^SPFp^ (**A**) and CGRP^PBel^ neurons (**B**). Scale bars indicate 100 μm. (**C**) Percentage of total projections from brain regions to CGRP^SPFp^ neurons. (**D**) Percentage of total projections from brain regions to CGRP^PBel^ neurons. (**E** and **F**) Diagram of the projection (%) from other brain regions to CGRP^SPFp^ (**E**) and CGRP^PBel^ neurons (**F**). CTX: cortex, Amy: amygdala, CP: striatum, GP: globus pallidus, BNST: bed nuclei of the stria terminalis, SI: substantia innominate, HPF: hippocampus, HY: hypothalamus, TH: thalamus, VTA: ventral tegmental area, SNr: substantia nigra, PRT: pretectal region, PAG: periaqueductal gray, SC: superior colliculus, IC: inferior colliculus, MRN: midbrain reticular nucleus, Pons: including the nucleus of the lateral lemniscus, pontine central gray, PBN, pontine reticular nucleus, VN: vestibular nucleus, NTS: nucleus tractus solitarius, SpV: trigeminal spinal nucleus, CBX: cerebellum.

**Figure S4.**
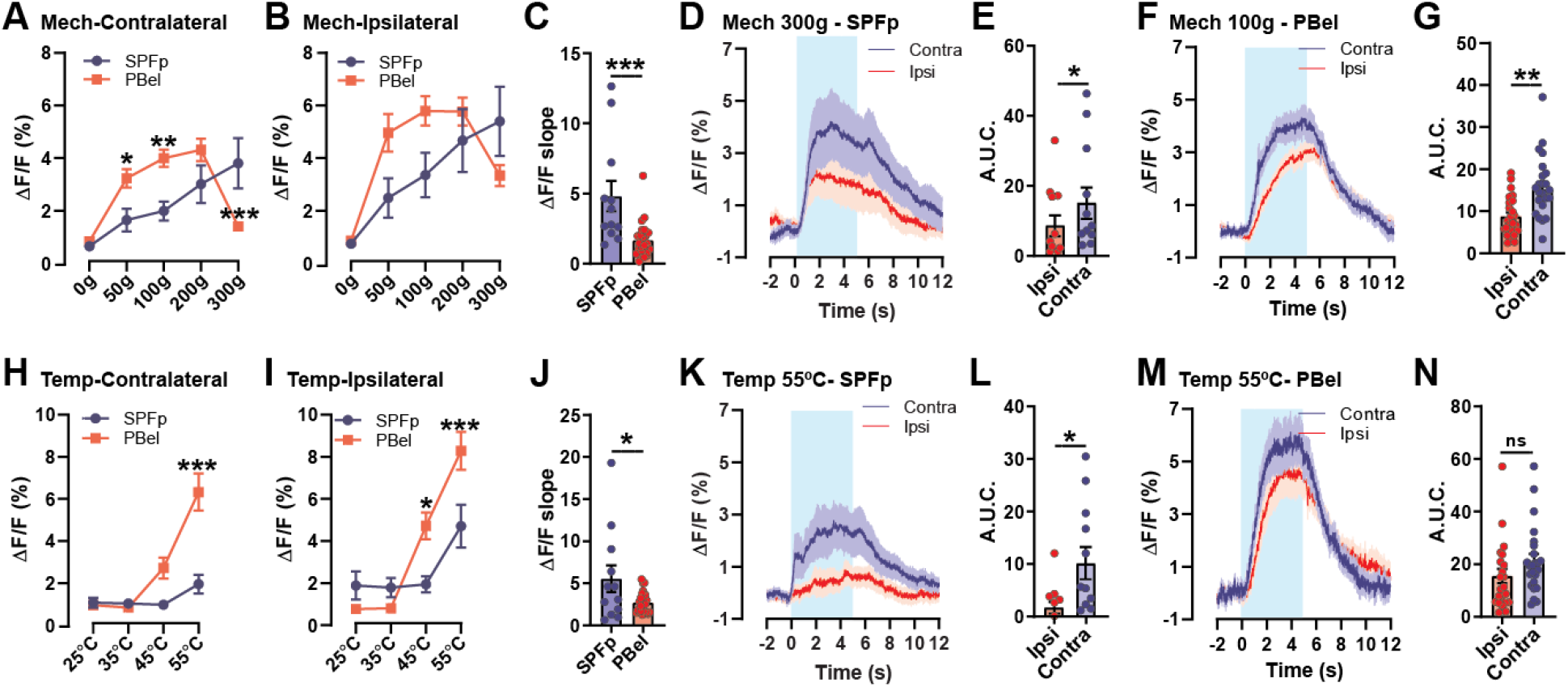
CGRP^SPFp^ and CGRP^PBel^ neurons are bilaterally stimulated by noxious stimuli. (**A and B**) Maximum calcium responses of the CGRP^SPFp^ and CGRP^PBel^ neurons to ipsilateral (**A**) and contralateral (**B**) mechanical stimulation. (**C**) CGRP^SPFp^ displayed faster increase of the calcium responses (bigger initial slope) to the contralateral mechanical stimulation (300 g for CGRP^SPFp^ and 100 g for CGRP^PBel^ neurons). (**D**-**G**) Calcium responses of CGRP^SPFp^ (**D** and **E**) neurons by ipsilateral or contralateral 300 g stimulation and CGRP^PBel^ (**F** and **G**) neurons by 100 g stimulation. (**H and I**) Maximum calcium responses of the CGRP^SPFp^ and CGRP^PBel^ neurons to ipsilateral (**H**) and contralateral (**I**) thermal stimulation. (**J**) Comparison of the initial slope of both neurons to contralateral 55°C stimulation. (**K**-**N**) Calcium responses of CGRP^SPFp^ (**K** and **L**) neurons and CGRP^PBel^ (**M** and **N**) neurons by ipsilateral or contralateral 55°C stimulation. **Statistics** (A) Repeated measure two-way ANOVA showed significance in intensity X region interaction (F(4, 136) = 11.80, p < 0.0001), intensity (F(4, 136) = 17.66, p < 0.0001), but not in region (F(1, 34) = 2.075, p = 0.1589). SPFp and PBel were significantly different in 50 (p < 0.05), 100 (p < 0.01) and 300 g (p < 0.001) with Sidak’s multiple comparisons test. (B) Repeated measure two-way ANOVA showed significance in intensity X region interaction (F(4, 136) = 5.468, p = 0.0004), intensity (F(4, 136) = 18.46, p < 0.0001), but not in region (F(1, 34) = 1.528, p = 0.2249). (C) SPFp: 4.82 ± 1.08 (n = 6 mice, 12 trial), PBel: 1.68 ± 0.26 (n = 6 mice, 24 trial). Unpaired t test (two-tailed), p = 0.0007. (E) SPFp; Ipsi: 8.51 ± 3.00 (n = 6 mice, 12 trial), Contra: 15.06 ± 4.51 (n = 6 mice, 12 trial). Paired t-test (two-tailed), p = 0.0498. (G) PBel; Ipsi: 8.79 ± 0.95 (n = 6 mice, 24 trial), Contra: 15.26 ± 1.61 (n = 6 mice, 24 trial). Paired t-test (two-tailed), p = 0.0012. (H) Repeated measure two-way ANOVA showed significance in intensity X region interaction (F(3, 102) = 8.995, p < 0.0001), intensity (F(3, 102) = 17.79, p < 0.0001), and region (F(1, 34) = 11.19, p = 0.002). SPFp and PBel were significantly different in 55 °C (p < 0.0001) with Sidak’s multiple comparisons test. (I) Repeated measure two-way ANOVA showed significance in intensity X region interaction (F(3, 102) = 10.16, p < 0.0001), intensity (F(3, 102) = 40.99, p < 0.0001), but not in region (F(1, 34) = 2.885, p = 0.0986). SPFp and PBel were significantly different in 45 (p < 0.05) and 55 °C (p < 0.0001) with Sidak’s multiple comparisons test. (J) SPFp: 5.55 ± 1.59 (n = 6 mice, 12 trial), PBel: 2.72 ± 0.26 (n = 6 mice, 24 trial). Unpaired t test (two-tailed), p = 0.0210. (L) SPFp; Ipsi: 1.74 ± 1.12 (n = 6 mice, 12 trial), Contra: 10.15 ± 3.08 (n = 6 mice, 12 trial). Paired t-test (two-tailed), p = 0.0146. (N) PBel; Ipsi: 15.55 ± 2.71 (n = 6 mice, 24 trial), Contra: 21.24 ± 2.63 (n = 6 mice, 24 trial). Paired t-test (two-tailed), p = 0.1385.

**Figure S5.**
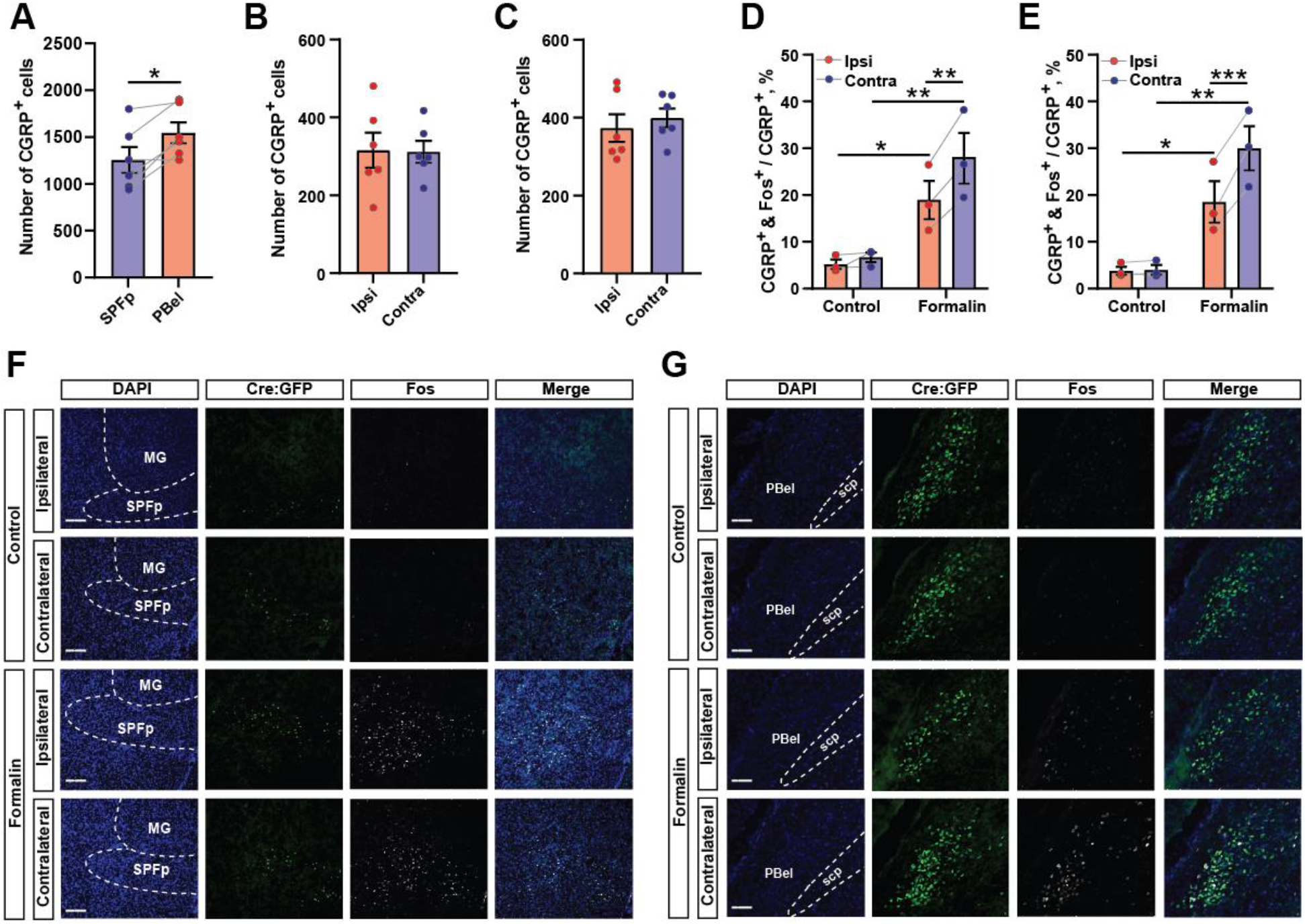
Bilateral activation of CGRP^SPFp^ and CGRP^PBel^ neurons by unilateral formalin injection in the same animal. (**A**) Number of CGRP positive cells in the SPFp and PBel. (**B** and **C**) The number of CGRP positive cells in each side of the SPFp (**B**) and PBel (**C**). (**D** and **E**) The percentage of CGRP neurons co-expressing c-Fos in the SPFp (**D**) and PBel (**E**). (**F** and **G**) Representative images of the SPFp (**F**) and PBel (**G**). Scale bars indicate 200 μm. **Statistics** (A) SPFp: 1255 ± 137.5, PBel 1545 ± 112.2 (n = 6 mice). Paired t-test (two-tailed), p = 0.0188. (B) SPFp; Ipsi: 315.5 ± 44.94, Contra: 311.8 ± 27.85 (n = 6 mice). Paired t-test (two-tailed), p = 0.9057. (C) PBel; Ipsi: 373.0 ± 35.34, Contra 399.3 ± 24.06 (n = 6 mice). Paired t-test (two-tailed), p = 0.2958. (D) SPFp; Repeated measure two-way ANOVA showed significance in treatment X side interaction (F(1, 4) = 17.42, p = 0.0140), treatment (F(1, 4) = 13.28, p = 0.0219), and side (F(1, 4) = 33.97, p = 0.0043). Ipsi vs contra was significant in formalin group (p < 0.01). Control vs formalin was significantly different in both ipsi (p < 0.05) and contra (p < 0.01) with Sidak’s multiple comparisons test. (E) PBel; Repeated measure two-way ANOVA showed significance in treatment X side interaction (F(1, 4) = 55.27, p = 0.0017), treatment (F(1, 4) = 19.57, p = 0.0115), and side (F(1, 4) = 58.66, p = 0.0016). Ipsi vs contra was significant in formalin group (p < 0.001). Control vs formalin was significantly different in both ipsi (p < 0.05) and contra (p < 0.01) with Sidak’s multiple comparisons test.

**Figure S6.**
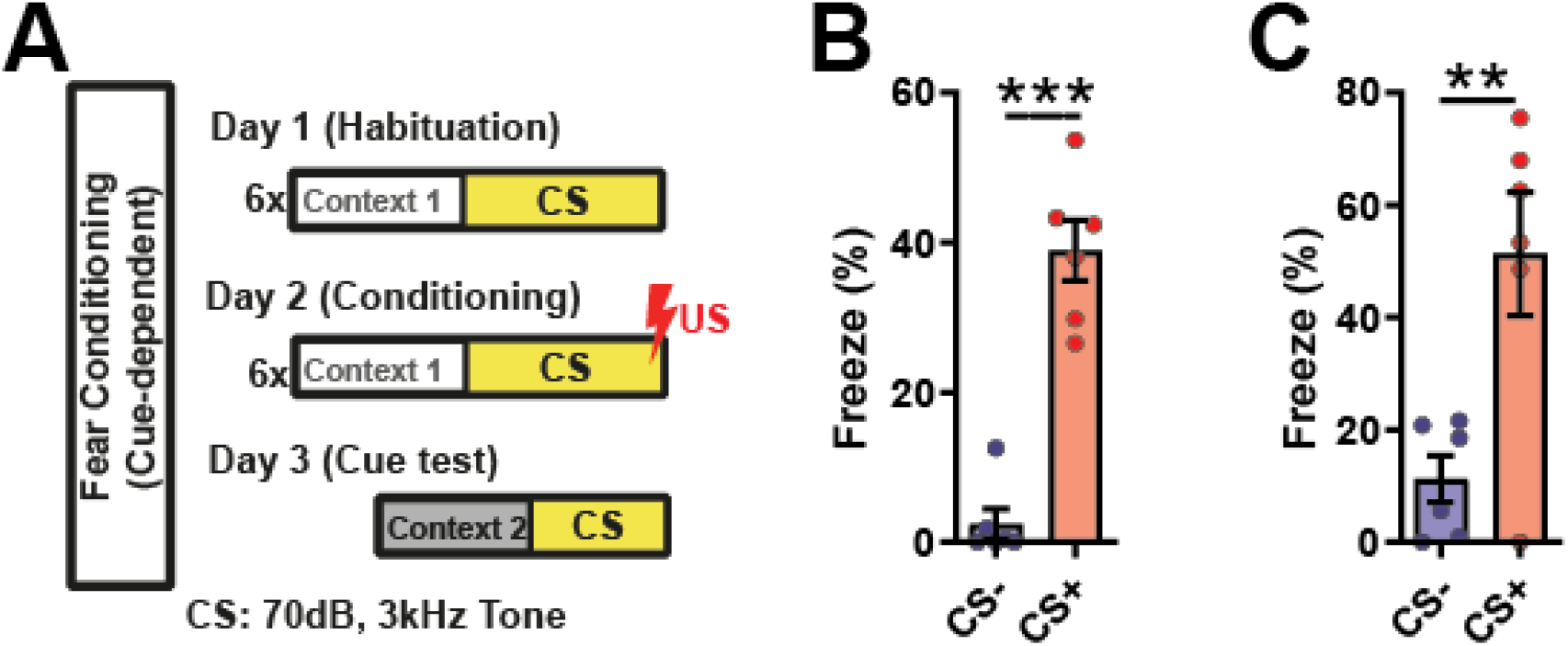
Cued fear conditioning with mice for CGRP^SPFp^ and CGRP^PBel^ fiber photometry. (**A**) Behavioral scheme for cued fear conditioning. Low intensity (70 dB, 3kHz) tone was used as CS in order not to induce calcium activity by sound. (**B** and **C**) Freezing was induced in both CGRP^SPFp^ (**B**) and CGRP^PBel^ (**C**) group. **Statistics** (B) SPFp; CS-: 2.47 ± 2.04 %, CS+: 38.96 ± 4.02 % (n = 6). Paired t test (two-tailed), p < 0.0001. (C) PBel; CS-: 11.27 ± 4.14 %, CS+ 51.31 ± 11.00 % (n = 6). Paired t test (two-tailed), p = 0.01.

**Figure S7.**
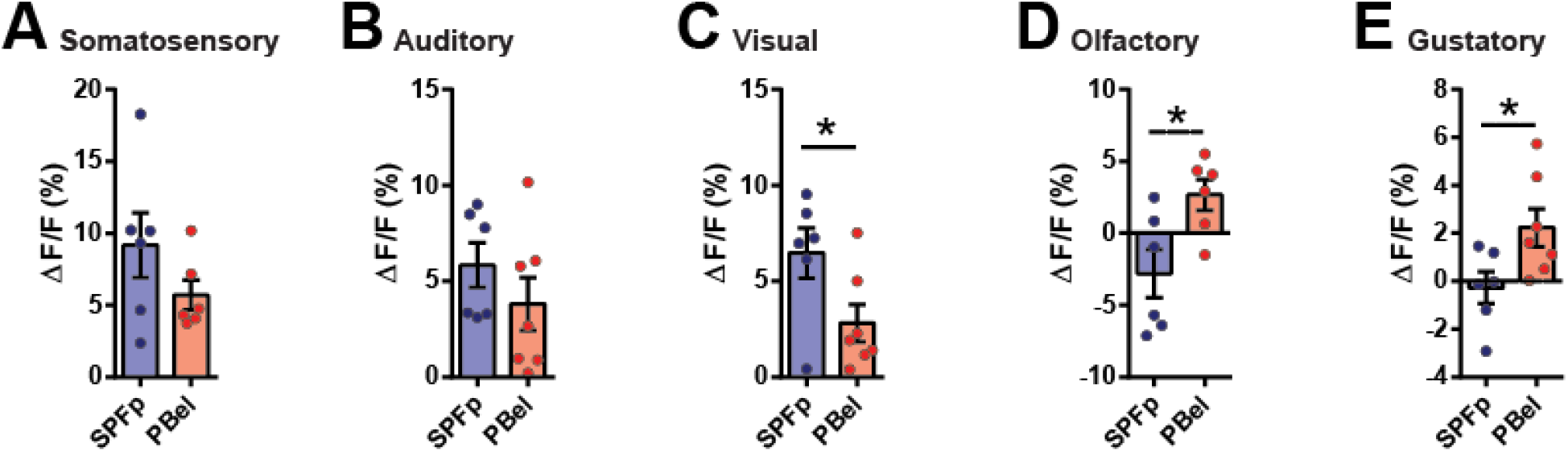
The CGRP^SPFp^ and CGRP^PBel^ neurons are differentially activated by multiple sensory threat cues. (**A**-**E**) Calcium response (peak amplitude) in the CGRP^SPFp^ and CGRP^PBel^ neurons by 2-s electric foot shock (0.6 mA) (**A**), 85-dB intense sound (**B**), rapidly expanding looming disk (**C**), TMT (**D**), and 0.5 mM quinine solution (**E**). **Statistics** (A) SPFp: 9.17 ± 2.25% (n = 6), PBel: 5.70 ± 1.03% (n = 6). Unpaired t test (two-tailed), p = 0.1917. (B) SPFp: 5.84 ± 1.17% (n = 6), PBel: 3.82 ± 1.39% (n = 7). Unpaired t test (two-tailed), p = 0.2987. (C) SPFp: 6.48 ± 1.31% (n = 6), PBel: 2.81 ± 0.96% (n = 7). Unpaired t test (two-tailed), p = 0.0418. (D) SPFp: −2.80 ± 1.68% (n = 6), PBel: 2.68 ± 1.07% (n = 6). Unpaired t test (two-tailed), p = 0.0206. (E) SPFp: −0.27 ± 0.66% (n = 6), PBel: 2.24 ± 0.79% (n = 7). Unpaired t test (two-tailed), p = 0.0363.

**Figure S8.**
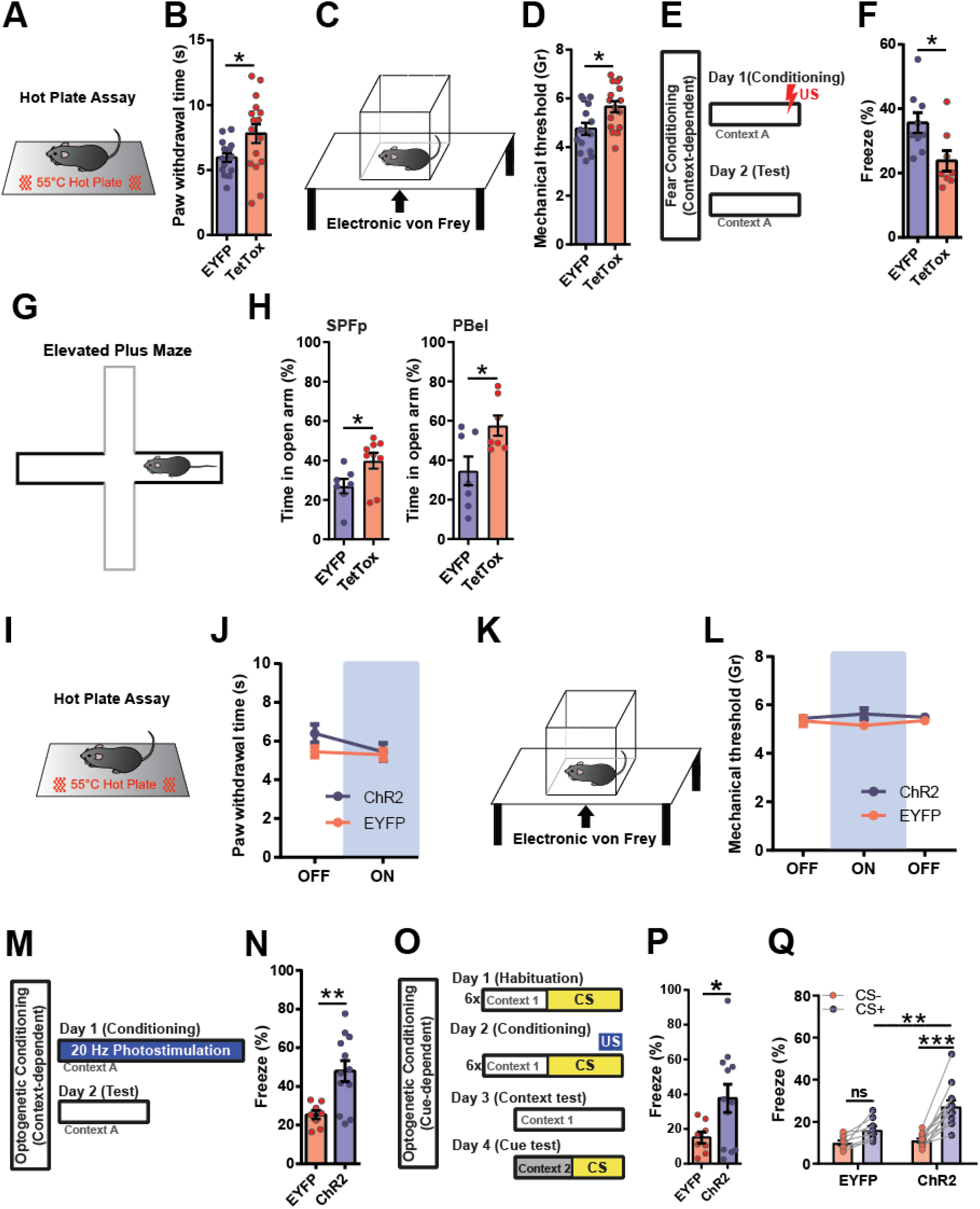
Manipulation of CGRP^SPFp^ neurons in behavior assays. (**A**, and **B**) Hot plate assay (55°C) with CGRP^SPFp^ silenced mice. (**C**, and **D**) Automatic von Frey assay with CGRP^SPFp^ silenced mice. (**E**) Experimental design for contextual fear conditioning. (**F**) Quantification of freezing 24 hr after contextual fear conditioning. (**G**) Schematic diagram of the elevated plus maze (EPM) test. (**H**) The EPM tests in CGRP^SPFp^ (left panel)-or CGRP^PBel^ neurons (right panel)-silenced mice. (**I**, and **J**) Optogenetic stimulation during hot plate assay (55°C). (**K**, and **L**) Optogenetic stimulation during electric von Frey assay. (**M**) Schematic diagram of optogenetic fear conditioning (photostimulation was used instead of foot shock). (**N**) Quantified freezing level 24 hr after optogenetic conditioning. (**O**) Schematic diagram of optogenetic cue-dependent fear conditioning (photostimulation paired with 3 kHz tone). (**P**) Context-dependent freezing 24 hr after optogenetic conditioning. (**Q**) Cue-dependent freezing 24 hr after optogenetic conditioning. **Statistics** (B) EYFP: 5.95 ± 0.33 s (n = 15), TetTox: 7.82 ± 0.73 s (n = 16). Unpaired t test (two-tailed), p = 0.0306 (D) EYFP: 4.75 ± 0.24 g (n = 15), TetTox: 5.66 ± 0.33 g (n = 16). Unpaired t test (two-tailed), p = 0.0113 (F) EYFP: 35.61 ± 3.18 % (n = 9), TetTox: 23.84 ± 3.17 % (n = 8). Unpaired t test (two-tailed), p = 0.0196 (H) SPFp; EYFP: 26.96 ± 3.66 % (n = 7), TetTox: 39.83 ± 4.05 % (n = 9). Unpaired t test (two-tailed), p = 0.0383. PBel; EYFP: 34.56 ± 7.37 % (n = 7), TetTox: 57.58 ± 5.15 % (n = 7). Unpaired t test (two-tailed), p = 0.0250. (I) Repeated measure two-way ANOVA showed no significance in interaction (F (1, 16) = 1.79, p = 0.2002), Laser (F (1, 16) = 3.36, p = 0.0857) and group (F (1, 16) = 1.49, p = 0.2393). (L) Repeated measure two-way ANOVA showed no significance in interaction (F (2, 36) = 1.72, p = 0.1938), Laser (F (2, 36) = 0.06, p = 0.9465) and group (F (1, 18) = 1.26, p = 0.2768). (N) EYFP: 25.31 ± 2.18 % (n = 8), ChR2: 47.95 ± 5.42 % (n = 12). Unpaired t test (two-tailed), p = 0.0042. (P) EYFP: 15.06 ± 3.20 (n = 8), ChR2: 37.55 ± 8.13 (n = 12). Unpaired t test (two-tailed), p = 0.044. (Q) Repeated measure two-way ANOVA showed significance in CS x group interaction (F (1, 18) = 8.072, p = 0.0108), CS (F (1, 18) = 39.57, p < 0.0001) and group (F (1, 18) = 6.827, p = 0.0176). Freezing at CS- and CS+ in ChR2 group (p < 0.0001) and CS+ in EYFP and ChR2 (p < 0.01) were significantly different with Sidak’s multiple comparisons test.

**Figure S9.**
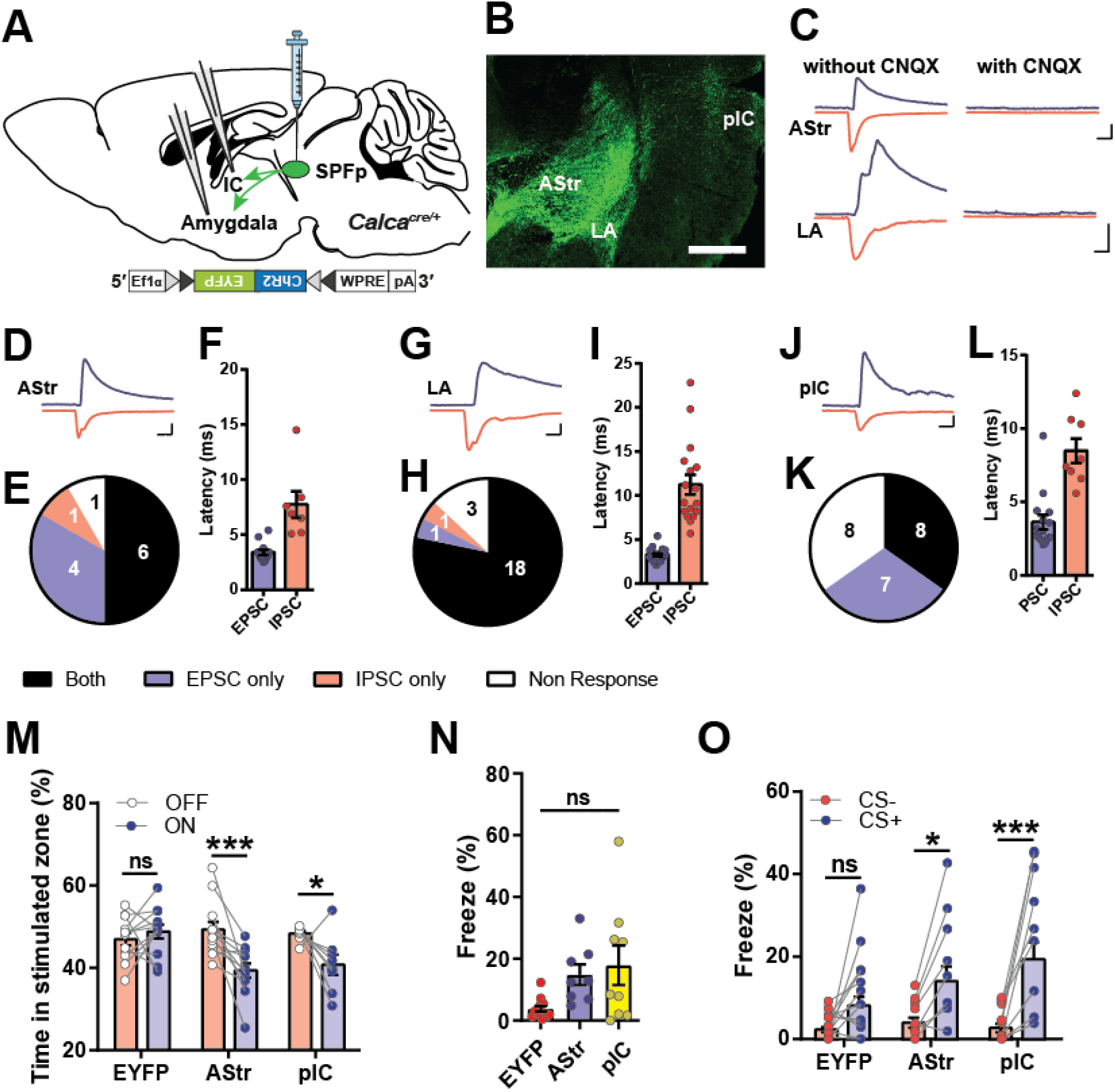
Mapping the functional downstream of CGRP^SPFp^ projection. (**A**) Schematics of the experiment. (**B**) Representative image of the projection regions from CGRP^SPFp^ neurons. Scale bar indicates 500 μm. (**C**) Example traces of an optically induced EPSC (blue) and IPSC (red) of Astr and LA without or with CNQX to confirm glutamatergic synapse. Scale bars indicate 10 ms and 50 pA. (**D**) Examples of Astr EPSC (blue) and IPSC (red) traces by optogenetic terminal activation. Scale bars indicate 10 ms and 50 pA. (**E**) Proportion of Astr cells with ‘Both’ EPSC and IPSC, ‘EPSC only’, ‘IPSC only’, or ‘No-Response’. (**F**) Onset of each Astr EPSC and IPSC to optogenetic stimulation. (**G**-**L**) Results of the same experiments with LA (**G**-**I**) and pIC (**J**-**L**) neurons. Scale bars indicate 10 ms and 50 pA. (**M**) Result of RTPA with terminal photo-stimulation. (**N**) Context-dependent freezing at 24 h after terminal photo-stimulation conditioning. (**O**) Cue-dependent freezing at 24 h after terminal photo-stimulation conditioning. **Statistics** (F) Astr; EPSC: 3.40 ± 0.25 ms (n = 12 cells), Astr IPSC: 7.74 ± 1.21 ms (n = 7 cells). Unpaired t test (two-tailed), p = 0.0003. (I) LA; EPSC: 3.30 ± 0.16 ms (n = 21 cells), LA IPSC: 11.24 ± 1.13 ms (n = 17 cells). Unpaired t test (two-tailed), p < 0.0001. (L) pIC; EPSC: 3.63 ± 0.49 ms (n = 15 cells), IC IPSC: 8.49 ± 0.83 ms (n = 8 cells). Unpaired t test (two-tailed), p < 0.0001. (M) Repeated measure two-way ANOVA showed significance in laser X group interaction (F(2, 29) = 7.215, p = 0.0029) and laser (F(1, 29) = 13.24, p = 0.0011, but not group (F(2, 29) = 2.607, p = 0.0910). Laser ON and was significantly larger than OFF in AStr (p < 0.001) and pIC (p < 0.05). Additionally, difference between EYFP vs Astr (p < 0.01) and EYFP vs pIC (p < 0.05) during ON period were significant. (N) EYFP: 3.842 ± 0.89 (n = 13), AStr: 14.82 ± 3.32 (n = 8), pIC: 17.94 ± 6.40 % (n = 9). One-way ANOVA showed significant (p = 0.0161). EYFP vs IC was significantly different (p < 0.05) with Tukey’s multiple comparison test. (O) Repeated measure two-way ANOVA showed significance in CS X group interaction (F(2, 27) = 3.647, p = 0.0396) and CS (F(1, 27) = 39.74, p < 0.0001) but not in group (F(2, 27) = 2.948, p = 0.0695). CS+ was significantly larger than CS-in AStr (p < 0.05) and pIC (p < 0.0001). Moreover, difference between EYFP and pIC in CS+ was significant (p < 0.01) with Sidak’s multiple comparison test.

**Figure S10.**
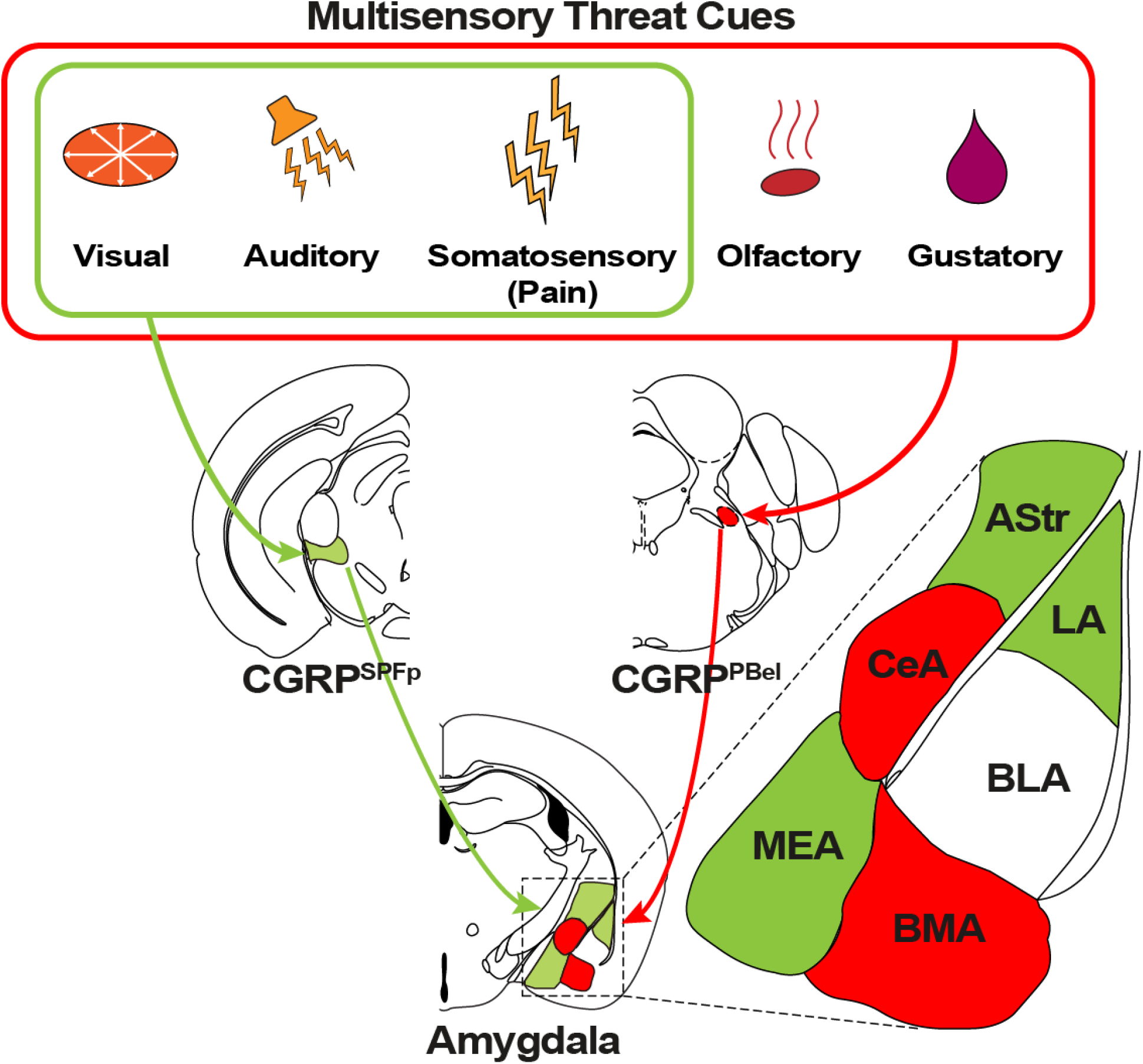
Summary illustration.

